# Cross-Species association statistics for genome-wide studies of host and parasite polymorphism data

**DOI:** 10.1101/726166

**Authors:** Hanna Märkle, Aurélien Tellier, Sona John

## Abstract

Uncovering the genes governing host-parasite coevolutionary interactions is of importance for disease management. The increasing availability of host and parasite full genome-data allows for cross-species genome-wide association studies based on the genomic data of infected hosts and their associated parasite strains sampled from natural populations (*i.e.* natural co-GWAs). Such studies focus on searching for cross-species allelic associations between pairs of host and parasite SNPs. We aim to quantify the power of natural co-GWAs to pinpoint loci under coevolution with respect to the intrinsic complexities of coevolutionary systems, such as the genetic specificity of the interaction and the temporal allele frequency changes resulting from the interaction. Therefore, we develop the cross-species association (CSA) and the cross-species prevalence (CSP) indices, the latter additionally incorporating genomic data from uninfected hosts. To provide an assessment of the statistical power of these indices, we analytically derive their genome-wide False Discovery Rates (FDR) based on the neutral site-frequency spectrum of the host and the parasite population. Using two coevolutionary models, we investigate under which parameter regimes these indices pin-point the coevolving loci. Under trench warfare dynamics, CSA and CSP are very accurate in pinpointing the loci under coevolution, while under arms race dynamics the power is limited especially for gene-for-gene interactions. Furthermore, we reveal that the combination of both indices across time samples is an indicator for the specificity of the interaction. Our results provide novel insights into the power and biological interpretation of natural cross-species association studies.

## 1. Introduction

The increasing availability of host and parasite whole-genome data provides powerful means to detect genes determining the outcome of host-parasite interactions. Recently, new Genome-Wide Association (GWA) methods to study host-parasite coevolutionary interactions have been proposed and performed (termed co-GWAs, Ebert 2018, MacPherson et al. 2018, Nuismer et al. 2017, Wang et al. 2018). The simple underlying idea is to compare all Single Nucleotide Polymorphisms (SNPs) between samples of hosts and parasites in a pairwise manner in order to detect significant correlations with the infection outcome (amount of infection, disease severity) or to detect significant allelex frequency correlations between species. However, such analyses rely on running large scale controlled experiments with numerous host and parasite genotypes (and thus can be termed controlled co-GWAs). A promising less-labor intensive alternative is to perform what we will call a natural co-GWAs. Such natural co-GWAs are based on sampling infected hosts and their associated parasites from natural populations (Ansari et al. 2017, Bartha et al. 2013, Bartoli and Roux 2017, Lees et al. 2019) and performing whole-genome sequencing of both interacting partners. The resulting data inherently contains phenotypic information (the infection outcome) about each sampled host-parasite pair, namely the given parasite genotype is infective on the given host genotype and hence, the host is susceptible to that particular parasite. The causal genetic variants for host susceptibility and parasite infectivity are expected to show statistically significant GxG associations (host genotype x parasite genotype) within the sample. A chief hypothesis is that these loci should explain a large proportion of the variance of the infection outcome as compared to loci in the neutral genomic background, which, by definition, have no effect on the infection outcome. To our knowledge, natural co-GWAs have been applied three times. These studies have uncovered strong associations between human SNPs 1) of the major histocompatibility complex (MHC) and known HIV epitopes (Bartha et al. 2013), 2) of leukocyte antigen molecules and components of the interferon lambda innate immune system and the hepatitis C virus NS5A protein (Ansari et al. 2017) and 3) of human disease susceptibility genes and *Streptococcus pneumoniae* genes responsible for invasiveness (Lees et al. 2019). Nevertheless, a key statistical issue, as in all GWAs, is that these natural co-GWAs methods rely on calculating association values for all possible comparisons of host and parasite SNP pairs, thus generating a distribution of millions of association values. As the top candidate loci are likely to be followed up on in molecular studies, there is a need to minimize the False Discovery Rate (FDR), that is the proportion of SNP pairs being identified as underlying the interaction when in fact they do not.

In principle, natural co-GWAs can be easily extended to any coevolutionary system where it is possible to call SNPs for a sample of infected hosts and the associated parasites (Bartoli and Roux 2017). An implicit underlying assumption is that the host and parasite genes revealed to determine the infection outcome are likely to be coevolving. Coevolution is defined as reciprocal changes in allele frequencies which result from selective pressures that two interacting species exert on one another (Janzen 1980). Allele frequency changes at coevolutionary loci are commonly described by a continuum between arms race (Dawkins and Krebs 1979, Stahl and Bishop 2000, Woolhouse et al. 2002) and trench warfare dynamics (Stahl et al. 1999). Arms race dynamics are characterized by recurrent fixation of alleles at the interacting loci and transient allelic polymorphism. Trench warfare dynamics, on the other extreme, result in long-term stable maintenance of several alleles with their frequencies either persistently fluctuating or converging towards a stable polymorphic equilibrium over time. Hence, allele frequency fluctuations are a common feature of all coevolutionary dynamics, irrespective of the ultimate fate of alleles. These allele frequency fluctuations are expected to affect the statistical power of natural co-GWAs over time. Indeed, it has been shown for experimental co-GWAS that the statistical significance of a given SNP pair association does depend on both host and parasite allele frequencies (MacPherson et al. 2018, Nuismer et al. 2017).

The speed and type of coevolutionary dynamics have been demonstrated to depend upon host and parasite life-history traits (reviewed in Brown and Tellier 2011), the strength of epidemiological dynamics (Ashby and Boots 2017) and the underlying host genotype by parasite genotype (G×G) interactions. Under the assumption that few major genes determine the interaction outcome, the underlying G×G interactions can be captured in a so called infection matrix 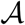. Here, each entry *α_ij_* denotes the probability that a parasite genotype *j* can infect a host genotype *i* (Tab. 1) or equivalently the degree of infection (disease severity). Two well known infection matrices are the matching-allele (MA) model and the gene-for-gene (GFG) model. In MA interactions a given parasite genotype can only infect a host when it matches the particular host allele (diagonal coefficients in Tab. 1b). For a 2×2 infection matrix the probabilities to infect the “non-matching” host genotypes can be defined as 1 − *ω*_1_ and 1 − *ω*_2_ (off diagonal coefficients in Tab. 1b). GFG interactions (Tab. 1c) are characterized by a universally susceptible host genotype (here host *i* = 1) and an universally infective parasite (here parasite *j* = 2). Here, the probability that host *i* = 2 is infected by parasite *j* = 1 is denoted by 1 − *ω*. Both matrices represent some points in a continuum of infection matrices (Agrawal and Lively 2002) and are a subset of more complex matrices with several alleles or loci (Ashby and Boots 2017, Gandon and Michalakis 2002).

**Table 1.** Infection matrices for coevolutionary models. The infection matrix 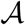 determines the outcome of the interaction between host genotypes (rows) and parasite genotypes (columns). Each *α_ij_* can be interpreted either as the probability for a given host to be infected or as the degree of infection (disease severity or partial resistance).

| a) general infection matrix | b) matching-allele | c) gene-for-gene |
| --- | --- | --- |
| $\begin{pmatrix} \alpha_{11} & \alpha_{12} \\ \alpha_{21} & \alpha_{22} \end{pmatrix}$ | $\begin{pmatrix} 1 & 1 - \omega_1 \\ 1 - \omega_2 & 1 \end{pmatrix}$ | $\begin{pmatrix} 1 & 1 \\ 1 - \omega & 1 \end{pmatrix}$ |

Our aim is to evaluate the statistical power and False Discovery Rate (FDR) of natural co-GWAs to pinpoint genes under coevolution given the various kinds of GxG interactions and the varying temporal dynamics of host-parasite coevolution. Therefore, we address the following three questions. First, which statistics (indices) are useful to locate the loci under coevolution? Second, what is the statistical power (FDR) of these indices under various infection matrices and/or fluctuating allele frequencies? Third, can these statistics provide some information about the underlying infection matrix and/or coevolutionary dynamics? To answer these questions, we first develop two indices: the cross-species association (CSA) index (analogous to the measure used in Ansari et al. 2017, Bartha et al. 2013 and suggested in Bartoli and Roux 2017) and the cross-species prevalence (CSP) index. Both indices measure the amount of association of allele frequencies across host and parasite loci. Second, we study six hypothetical numerical examples representing varying amounts of cross-species association. For these examples, we calculate the introduced indices along with p-values of a corresponding *χ*^2^-test. The p-value allows us to measure the chance that a pair of loci under coevolution would be among the top candidates when correcting for multiple testing. Third, we explicitly calculate the expected distribution of the values of both indices for all possible comparisons between pairs of neutral host and neutral parasite loci (representing the genomic background). Our computations are based on the site-frequency spectrum distribution of neutral allele frequencies for a population with constant size. This neutral distribution of values of our indices allows us to access the False Discovery Rate (FDR), that is the percentage of pairs of neutral alleles which show higher index values than the pair of loci under coevolution. We can show for these same hypothetical examples that our FDR results are in accordance with those of the *χ*^2^-test. Fourth, we present analytical results on the expected values of our indices at the loci under coevolution for two widely used coevolutionary models and for arbitrary infection matrices. Finally, we quantify the statistical power of these statistics to pinpoint coevolving loci over the course of a coevolutionary interaction and for different underlying GxG matrices by terms of simulation. To do so, we calculate both indices over the entire course of a coevolutionary dynamic and access the FDRs based on the neutral distributions. We demonstrate that performing co-GWAs using these indices across time samples not only allows to infer 1) the type of dynamics occurring (arms race versus trench warfare), but also 2) the underlying GxG interaction matrix. We conclude by discussing the applicability of our co-GWAs indices to studies of coevolution in natural or controlled systems.

## 2. Methods

### 2.1. Definition of the association indices

We assume that *n*_T_ host individuals are sampled and genotyped at each biallelic single nucleotide polymorphism (SNP) in the genome, so that at any locus *l*, two types of hosts are found (*i* ∈ (1, 2)). The total host sample *n*_T_ consists of *n*_Inf_ infected hosts, the infected subsample, and *n*_H_ non-infected (healthy) hosts, the non-infected subsample. A number of *n*_Par_ parasite samples is obtained from the *n*_Inf_ infected hosts (one sample per host) and also genotyped at each biallelic SNP. Accordingly, there are also two parasite types for each biallelic SNP (*j* ∈ (1, 2)). Note that bi-allelic SNPs are also a common assumption in GWAs and co-GWAs. Our definition also applies to any type of mutation with two states such as insertion-deletion (of few to many base pairs) or presence/absence polymorphism of larger genomic regions (*e.g.* coding genes or transposable elements).

#### 2.1.1. The Cross-Species Association index (CSA)

We define the absolute Cross Species Association index (CSA) when sampling *n*_Inf_ hosts and *n*_Par_ (= *n*_Inf_) parasites as:

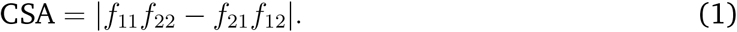

Here, *f_ij_* is the proportion of hosts of type *i* being infected by a parasite of type *j* in the infected subsample (*n*_Inf_), so that 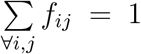. The statistic is an adaptation of the well-known linkage disequilibrium (LD) measure in population genetics (Charlesworth and Charlesworth 2010, Lewontin and Kojima 1960, p371-373).

Following population genetics theory (Charlesworth and Charlesworth 2010, p371-373), we normalize CSA in two different ways such that the absolute values range from 0 to 1. First, we define CSA′ by normalizing CSA values by the maximum CSA value possible, CSA^max^ = 0.25.

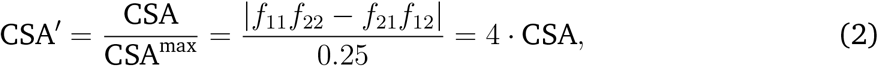

CSA is at its maximum when hosts of type 1 are solely infected by parasites of type 1 and hosts of type 2 are solely infected by parasites of type 2 and *f*_11_ = *f*_22_ = 0.5 (or when hosts of type 1 are solely infected by parasites of type 2 and hosts of type 2 are solely infected by parasites of type 1 and *f*_12_ = *f*_21_ = 0.5).

The second normalization consists in dividing the CSA value by the square root of the product of the frequencies of the different host and parasite alleles in the infected subsample

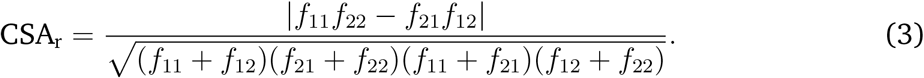

#### 2.1.2. The Cross-Species Prevalence index (CSP)

We define the Cross Species Prevalence index (CSP) as

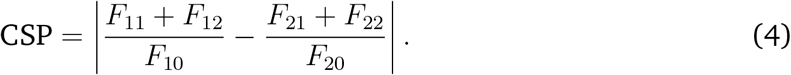

Here, *F_ij_* is the proportion of host type *i* infected by parasite type *j* in the total host sample (*n*_T_) and *F_i_*_0_ is the proportion of uninfected hosts of type *i* in the total sample (*n*_T_). By definition, *F*_11_ + *F*_12_ + *F*_21_ + *F*_22_ = *n*_Inf_*/n_T_*, *F*_10_ + *F*_20_ = (*n_T_* − *n*_Inf_)*/n_T_*, and *F*_11_ + *F*_12_ + *F*_21_ + *F*_22_ + *F*_10_ + *F*_20_ = 1 (see Fig. 1). Note that *F_i_*_0_ is composed of individuals which either 1) have no interaction with any parasite if the disease prevalence is less than 100%, or 2) are exposed to parasites but resist infection. CSP is defined as long as there are uninfected individuals of both host types present in the population (all *F_i_*_0_ ≠ 0). CSP is thus an indirect association index as it does not explicitly take into account which parasite type infects which host type in the infected subsample. But, it depends on the frequencies of the host types in the different subsamples which are determined by the specificity of the interaction and the parasite allele frequencies.

**Figure 1.**
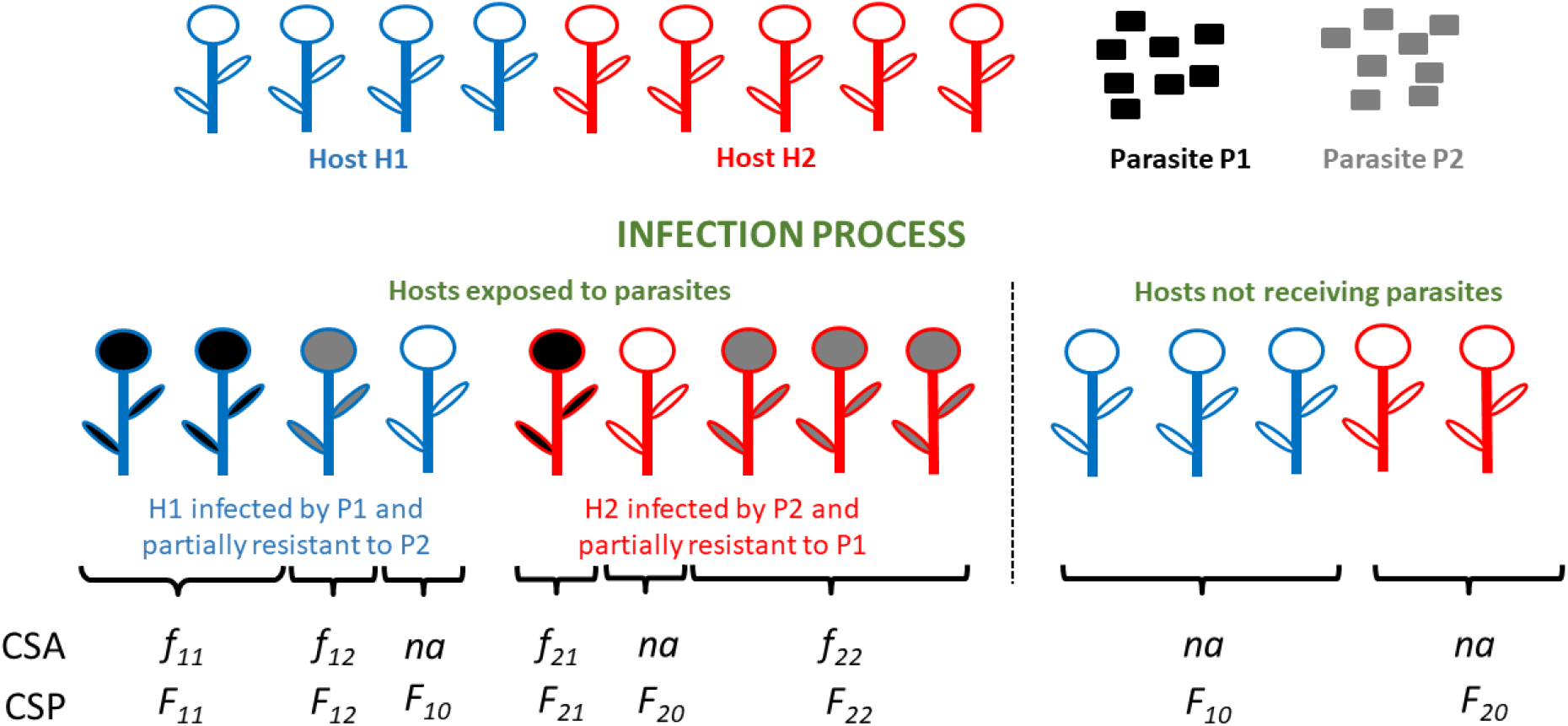
Schematic representation of the CSA and CSP definitions. The host population consists of two host types *H*_1_ (square) and *H*_2_ (circle) and the parasite population consists of two parasite types *P*_1_ (black) and *P*_2_ (grey). A proportion *ϕ* of the hosts is exposed to parasites. Hosts exposed to the parasite either become infected or they can resist infection. Infected hosts are colored based on the identity of the infecting parasite genotype (grey or black). *f_ij_* is the proportion of hosts type *i* infected by parasites of type *j* among infected hosts. *F_ij_* is the proportion of hosts of type *i* infected by parasites of type *j* in the whole host population (sum of all hosts). *F_i_*_0_ is the proportion of non-infected hosts of type *i* in the whole host population. *F_i_*_0_ is composed of type *i* hosts which did not receive spores (1 − *ϕ*) or received spores and are resistant to the respective parasite.

### 2.2. Statistics, FDR and distributions of indices’ values

When performing a natural co-GWAS among *m_H_* host SNPs and *m_P_* parasite SNPs, there is a potentially very large total number of *m_H_* ∗ *m_P_* comparisons (see Ansari et al. 2017, Bartha et al. 2013). This means that any significance threshold based on neutral SNPs should be stringent enough to have a low number of false candidate loci (a low FDR).

#### 2.2.1. *χ*^2^-test for CSA/CSP

For both CSA and CSP indices, we can compare proportions in a 2×2 contingency table, and thus perform a simple *χ*^2^-test of homogeneity to obtain the p-value for a given pair of SNPs. The corresponding contingency table for CSA is

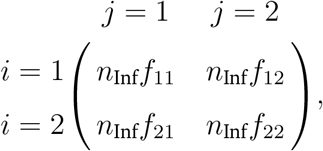

and the corresponding contingency table for CSP is

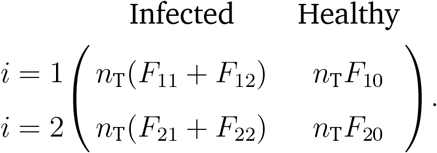

To account for multiple testing, we apply a Bonferroni-correction by dividing the chosen significance threshold (*e.g.* 0.05 or 0.01) by the total number of comparisons (*m_H_* ∗ *m_P_*). We provide R-codes to compute the values of CSA and CSP and corresponding *χ*^2^-tests for a list of pairwise comparisons between *m_H_* host and *m_P_* parasite SNPs.

#### 2.2.2. Expected distributions of CSA and CSP for neutral SNPs

To quantify the statistical power (FDR) of our indices without having empirical data at hand, we compute the expected theoretical distribution of CSA and CSP values based on a large number of pairs of neutral host and parasite loci. Full genome data usually contains a large number of neutral SNPs which are distributed across the whole genome. Given that the recombination rate is high enough, these neutral SNPs are assumed to evolve independently from one another and thus, both their population and sample allele frequencies are mutually independent. Therefore, we can conveniently draw the sample frequencies for our neutral loci from an expected neutral site-frequency spectrum for a population with constant size.

By definition, neutral host and parasite loci do not determine the outcome of infection. Therefore, each neutral hostt SNP-neutral parasite SNP pair is characterized by an infection matrix 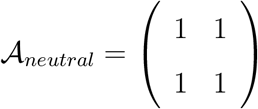.

We obtain the expected value for CSA as follows:

1. We calculate the expected value of CSA when a host SNP has minor allele frequency *v* and a parasite SNP has minor allele frequency *w* in the infected subsample *n*_Inf_. Given these minor allele frequencies, we calculate the value of CSA for each possible configuration of alleles and weight it by the probability of the configuration.

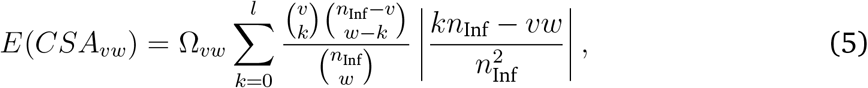

where *l* = *min*(*v, w*), and Ω_*vw*_ is the normalization for either obtaining CSA′ (with Ω_*vw*_ = 4, from Eq. 2) or CSA_*r*_ (with_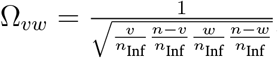_, from Eq. 3).
2. Based on the neutral host and parasite site frequency spectra we calculate the probability *p_v_* that a host SNP has minor allele frequency *v* and the probability *p_w_* that a parasite SNP has minor allele frequency *w* in a sample of size *n*_Inf_.
3. We weight the expected values obtained in step (1) by the probabilities obtained in step (2)

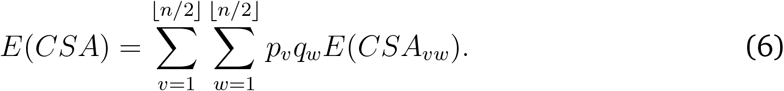

This approach also allows us to generate a probability mass function and a cumulative distribution function for a given sample size, from which the FDR is computed by using the r-package distr (Ruckdeschel et al. 2006).

For the CSP we proceed in a similar way.

1. We calculate the expected value of CSP when a host SNP has minor allele frequency *v* in the total host sample *n*_T_ and a parasite SNP has minor allele frequency *w* in the infected subsample *n*_Inf_. In contrast to the CSA calculations, we first h ave to access all possible ways of assigning host alleles to the healthy (*n*_H_) and infected (*n*_Inf_) subsample. We restrict ourselves to configurations where both alleles can be found in both the infected and the healthy subsample.

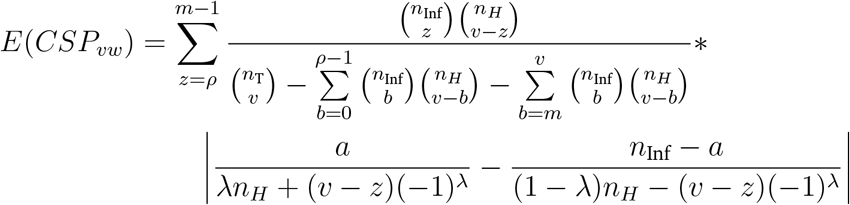

with *ρ* = *max*(1*, v* − *n_H_* + 1), *m* = *min*(*n*_Inf_*, v*), *a* = *min*(*z, n*_Inf_ − *z*), and

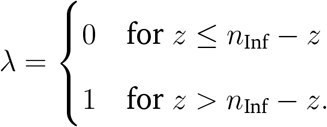
2. Based on the neutral host and parasite site frequency spectra, we calculate the probability *p_v_* that a host SNP has minor allele frequency *v* in a sample of size *n*_T_ and the probability *p_w_* that a parasite SNP has minor allele frequency *w* in a sample of size *n*_Inf_.
3. We weight the expected values obtained in step (1) by the probabilities obtained in step (2)

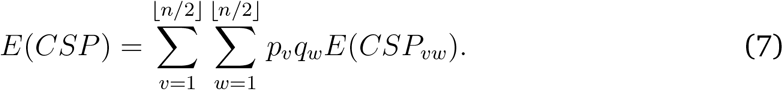

Without loss of generality, we use the folded SFS of both the host and the parasite. Thus, we assume that there is no outgroup sequence available and the ancestral and derived state are indistinguishable at each SNP. For simplicity, we assume that the allele frequency distribution of neutral SNPs in both, the host and the parasite, follows the SFS under drift-mutation equilibrium for a Wright-Fisher model under constant population size (our Eq. S2 is found for example in Durrett 2010, p.50). We compute the distributions for CSA and CSP for different sample sizes and sampling schemes. For any CSA or CSP value at a pair of coevolving loci, we can extract the False Discovery Rate (FDR) from this distribution assuming that there is only one or very few coevolving pairs of loci. The FDR is the proportion of pairs of neutral loci which exhibit a higher index value than a pair of coevolving loci. Based on the design of realistic molecular biology studies, we aim to obtain a maximum of ten candidate pairs of loci. Keeping in mind that both indices are computed for *m_H_* ∗ *m_P_* pairs of loci, the FDR rate therefore must be roughly smaller than 10*/*(*m_H_* ∗ *m_P_*). If not stated otherwise, we show results for *n_T_* = 200 and *n*_Inf_ = *n*_H_ = 100 (see the Online Supplementary Material for other sampling schemes Fig. S1 - S5).

### 2.3. The coevolutionary models

We study the temporal dynamics of CSA and CSP at a pair of coevolving loci using two different models, namely a pure population genetics model and an epidemiological model which incorporates eco-evolutionary feedback. Both models can be flexibly parameterized in terms of the underlying infection matrix and have been shown to exhibit the full range of coevolutionary dynamics.

#### 2.3.1. Model A: population genetics model

First, we use a simple population genetics model (henceforward termed model A) to study the allele frequency changes at the coevolving loci under the assumption of very large (infinite) haploid host and parasite population sizes. We assume that host and parasite generations are discrete and synchronized in terms of reproduction (Tellier and Brown 2007b). The frequency of host genotype *h_i_* and parasite genotype *p_j_* in generation *g* + 1 is obtained as:

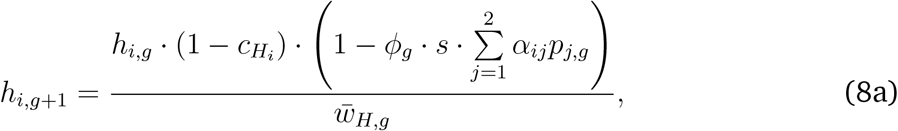

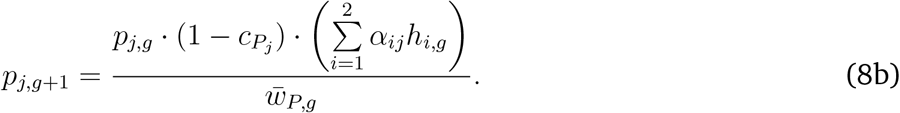

Every generation *g*, a proportion *ϕ_g_* of the host population (*i.e.* the exposed fraction) interacts with the parasite population in a frequency-dependent manner. The outcome of interactions is determined by the infection matrix 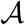. Therefore, the proportion of infected individuals in the population (*i.e.* the disease prevalence) is 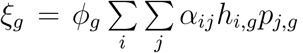. An infection reduces the relative fitness of hosts by an amount *s* (cost of infection). Further, each host genotype *i* (parasite genotype *j*) can be associated with some fitness cost 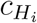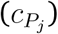, such as a cost of resistance (infectivity). Unlike the cost of infection (*s*) which reduces the fitness of only the infected individuals, the cost of resistance 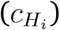 and cost of infectivity 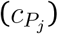 reduce the fitness of all individuals of the given genotype, irrespective of the infection status. Previous work has shown that this model always generates arms race dynamics (Tellier and Brown 2007b).

This dynamical system admits an equilibrium point when the conditions *h_i,g_*_+1_ = *h_i,g_* = *ĥ_i_* and 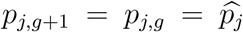 hold for each host genotype *i* and each parasite genotype *j*. There are four so called trivial monomorphic equilibrium points at which one host and one parasite allele are fixed, and one polymorphic equilibrium with frequencies:

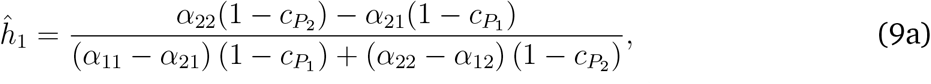

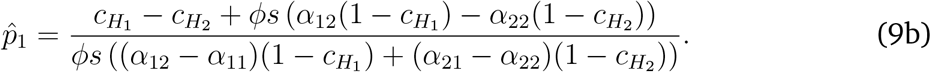

Under this model, the value of CSA at generation *g* (Eq. 1) can be obtained as:

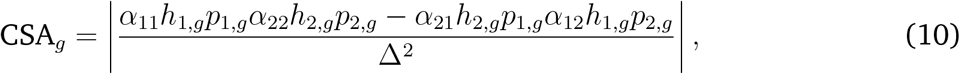

where Δ = *α*_11_*h*_1,*g*_*p*_1,*g*_ + *α*_22_*h*_2,*g*_*p*_2,*g*_ + *α*_21_*h*_2,*g*_*p*_1,*g*_ + *α*_12_*h*_1,*g*_*p*_2,*g*_ (introduced to make sure in Eq. 1 that 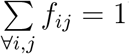.

When Eq. 4 is applied to our coevolutionary model A, the CSP value at each generation *g* is obtained as:

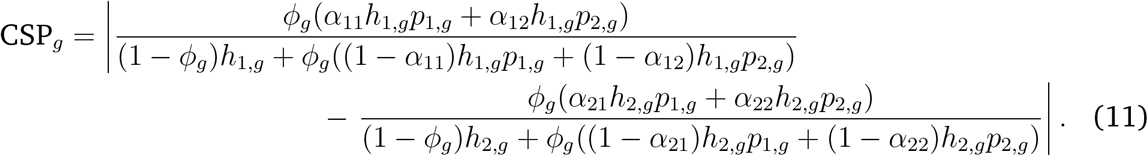

Based on these equations we can calculate the value of CSA and CSP at the non-trivial equilibrium point under both a MA and a GFG infection matrix.

#### 2.3.2. Model B: model with epidemiological dynamics and feedback

In model B we consider a continuous time coevolutionary model (Živković et al. 2019) based on a known Susceptible-Infected model (Ashby and Boots 2017, Boots et al. 2014, May and Anderson 1983). This model allows for simultaneous changes in population sizes and allele frequencies. The total number of hosts of type *i* includes *S_i_* susceptible and Σ*_j_ I_ij_* infected individuals. The change in number of susceptible hosts *S_i_* is given by Eq. 12a and the change in number of infected individuals *I_ij_* is given by Eq. 12b.

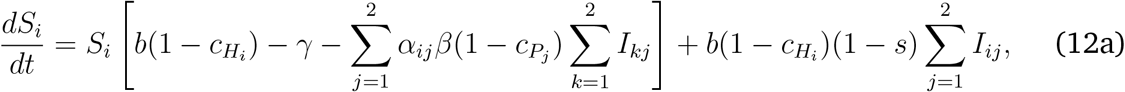

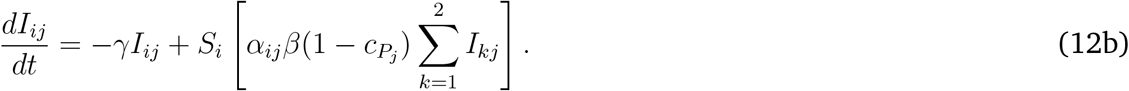

The number of parasites of type *j* is obtained as 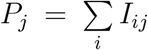 and hence, the change in number of parasites of type *j* is given by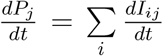. Hosts reproduce at natural birth rate *b* and die at natural death rate *γ*. The total host population size at generation *t* is 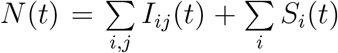. We assume that there is no vertical disease transmission, and the infections are sustained in the population through an overlap between generations. Uninfected hosts can get infected by horizontal disease transmission at rate *β*. The costs 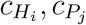 and *s* are defined as in model A. Previous analyses have shown that, depending on the parametrization (chosen infection matrix and parameter values), this model results in a range of different dynamics (arms race dynamics, trench warfare dynamics with stable limit cycles and trench warfare dynamics with a stable attractor) (Ashby and Boots 2017, Živković et al. 2019). We focus here chiefly on the trench warfare outcome, and especially when the dynamics converge to a stable (attractor) polymorphic equilibrium point.

To simulate the dynamics, we discretize model B into small time steps of size *δ_t_*. Hence, one discrete generation *t* consists of 1*/δ_t_* time steps. The value of *δ_t_* is chosen so that the discretized time dynamics match the continuous time dynamics. The changes in population sizes and in allele frequencies, as well as the corresponding association statistics values, are computed over time and at the equilibrium point. The disease prevalence is here an inherent property of the disease dynamics and allele frequencies, as defined under the eco-evolutionary feedbacks (Ashby et al. 2019, Boots et al. 2014), and thus varies over time:

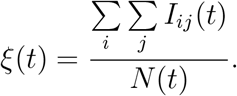

For Model B, the CSA value at each time *t* is obtained as:

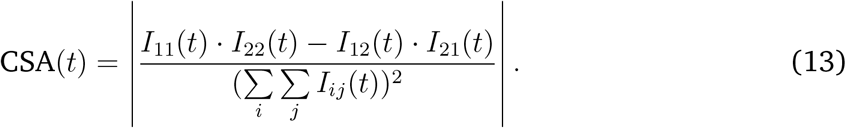

For Model B, the CSP value at time *t* is given by:

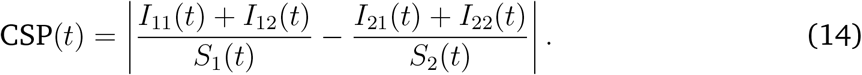

#### 2.3.3. Numerical simulations

For both models we conduct numerical simulations under a symmetric matching-alleles (Tab. 1a), an asymmetric matching-alleles (Tab. 1b) and a GFG-interaction (Tab. 1c) interaction over 500 generations. In line with previous studies we assume no genotype costs 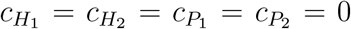 for both symmetric and asymmetric MA interactions (Gandon and Nuismer 2009). Under a GFG interaction model we assume that 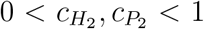 and 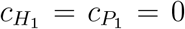 (Tellier and Brown 2007b). We calculate the values of CSA_*r*_/CSA′/CSP at any generation, based on the equations above (Eqs. 10, 11, 13, 14) and compare them to a FDR of 1%, based on the theoretical SFS distributions for neutral loci computed in section 2.2.2.

We also access the power of using time sample data to pinpoint the coevolutionary locus. As in natural populations it may prove difficult to obtain time samples spanning extended periods of time, we use a simple, realistic and affordable (in terms of sequencing costs) scheme with three time samples taken within an interval of ten generations. In natural systems the state of coevolutionary dynamics is usually unknown, and thus on purpose, we do not optimize the sampling strategy to detect coevolutionary cycles. We present the analysis for two sampling schemes, denoted *Z*_1_ and *Z*_2_, which are chosen to represent the early and later phase of the simulations. For all dynamics, we fix *Z*_1_ to consist of time samples from generations 50, 55 and 60 and *Z*_2_ from generations 250, 255 and 260. All codes used for generating and analyzing the data can be obtained from: URL XY.

## 3. Results

### 3.1. Hypothetical numerical examples

In order to explicit the rationale behind our indices, we consider six cases with decreasing levels of association (maximum association for case 1, random association for case 6). Case 1 for CSP in Tab. 3 is chosen such that *F_i_*_0_ ≠ 0 and therefore, CSP is defined. We compute the p-value of a *χ*^2^-test for each case. Both our indices reflect the strength of the association, as the FDR is very low for the example with the strongest association, especially for CSA′ (Table 2, Table 3). Further, we show that our indices and their FDR values show the same trend as the p-value of the corresponding *χ*^2^-test. When correcting for multiple testing by applying a Bonferroni-correction based on a arbitrary (but realistic, see Ansari et al. 2017, Bartha et al. 2013) number of *m_H_* ∗ *m_P_* = 10^7^ comparisons, the SNP-pairs of cases 1, 2 and 3 in Tab. 2 would be still significant under the *χ*^2^-test. However, the FDR CSA_r_ for case 2 is around 4*e*^−4^, which would result in 4, 000 false positive SNP pairs under *m_H_* ∗ *m_P_* = 10^7^. In contrast, the FDR for CSA′ is lower than 10^−12^, which would correspondingly translate into only a handful (if any) false positive SNP pairs. Nevertheless, our indices are robust and stringent (low FDR) as they will only pinpoint with high accuracy the host-parasite SNP-pair, which represents the highest level of association.

**Table 2.** Six hypothetical numerical examples to illustrate the strength of association when sampling only infected individuals and the relevant statistics: the p-value of a *χ*^2^-test, CSA value and the corresponding false discovery rate (FDR) based on neutral allele frequencies. The examples are build assuming both host and parasite loci with a minor allele frequency of 0.5. The FDR is calculated for a sample size of *n*_Inf_ = 100. Asterisks (*) indicate significant p-values after Bonferroni correction for 10^7^ comparisons.

|  | Case 1 | Case 2 | Case 3 | Case 4 | Case 5 | Case 6 |
| --- | --- | --- | --- | --- | --- | --- |
| $f_{11}$ | 0.5 | 0.45 | 0.40 | 0.35 | 0.30 | 0.25 |
| $f_{12}$ | 0 | 0.05 | 0.10 | 0.15 | 0.20 | 0.25 |
| $f_{21}$ | 0 | 0.05 | 0.10 | 0.15 | 0.20 | 0.25 |
| $f_{22}$ | 0.5 | 0.45 | 0.40 | 0.35 | 0.30 | 0.25 |
| p-value $\chi^2$ | $1.52e^{-23*}$ | $1.24e^{-15*}$ | $1.97e^{-9*}$ | $6.33e^{-5}$ | $4.55e^{-2}$ | 1 |
| CSA value | 1 | 0.8 | 0.6 | 0.4 | 0.2 | 0 |
| FDR CSA<sub>r</sub> | $< 10^{-12}$ | $3.9e^{-4}$ | $1.2e^{-3}$ | $4.5e^{-3}$ | $4.8e^{-2}$ | 1 |
| FDR CSA' | $< 10^{-12}$ | $< 10^{-12}$ | $3.04e^{-12}$ | $6.81e^{-7}$ | $1.88e^{-3}$ | 0.996 |

**Table 3.** Six hypothetical numerical examples to illustrate the strength of association when sampling infected and non-infected individuals and the relevant statistics: the p-value of a *χ*^2^-test, CSP value and the corresponding false discovery rate (FDR) based on neutral allele frequencies. The examples are build assuming both host and parasite loci with a minor allele frequency of 0.5. The FDR is calculated for sample sizes *n_T_* = 200 and *n*_Inf_ = 100. Asterisks (*) indicate significant p-values after Bonferroni correction for 10^7^ comparisons.

|  | Case 1 | Case 2 | Case 3 | Case 4 | Case 5 | Case 6 |
| --- | --- | --- | --- | --- | --- | --- |
| $F_{11} + F_{12}$ | 0.49 | 0.45 | 0.40 | 0.35 | 0.30 | 0.25 |
| $F_{10}$ | 0.01 | 0.05 | 0.10 | 0.15 | 0.20 | 0.25 |
| $F_{21} + F_{22}$ | 0.01 | 0.05 | 0.10 | 0.15 | 0.20 | 0.25 |
| $F_{20}$ | 0.49 | 0.45 | 0.40 | 0.35 | 0.30 | 0.25 |
| p-value $\chi^2$ | $5.52e^{-42*}$ | $1.12e^{-29*}$ | $2.15e^{-17*}$ | $1.54e^{-8}$ | $4.68e^{-3}$ | 1 |
| CSP value | 48.98 | 8.889 | 3.75 | 1.905 | 0.833 | 0 |
| FDR CSP | $< 10^{-12}$ | $1.69e^{-4}$ | $7.1e^{-3}$ | $3.83e^{-2}$ | 0.12 | 0.8044 |

### 3.2. Analytical results for model A

In the methods and material section we computed the threshold for detecting pairs of coevolving SNPs for a large number of pairwise SNP comparisons. We now quantify the usefulness and power of our statistics to uncover genes under coevolution in a natural setting with coevolutionary dynamics and different underlying infection matrices.

#### 3.2.1. Under a matching-alleles infection matrix

For a matching-allele infection matrix (Table 1b) and for 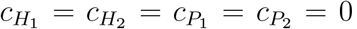, the frequencies at the non-trivial equilibrium point in model A (Eq. 9) are:

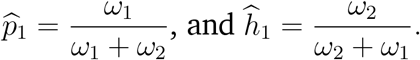

Combining these equilibrium frequencies with Eq. 10 and Eq. 11 we can obtain the values of the indices at the polymorphic equilibrium point.

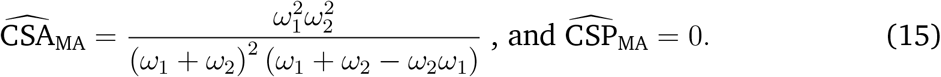

For a matching alleles (MA) interaction without any genotype costs, CSP is always zero at the equilibrium point (Eq. 15), irrespective of the values of *ω*_1_ and *ω*_2_. Under the MA model, the values of the CSA and CSP display a different behavior over time and the value of CSA at the equilibrium depends on *ω*_1_ and *ω*_2_. This implies that tracking CSA and CSP values over time yields insights into the asymmetry of the infection matrix (Eq. 15, Figs. 2, 4).

**Figure 2.**
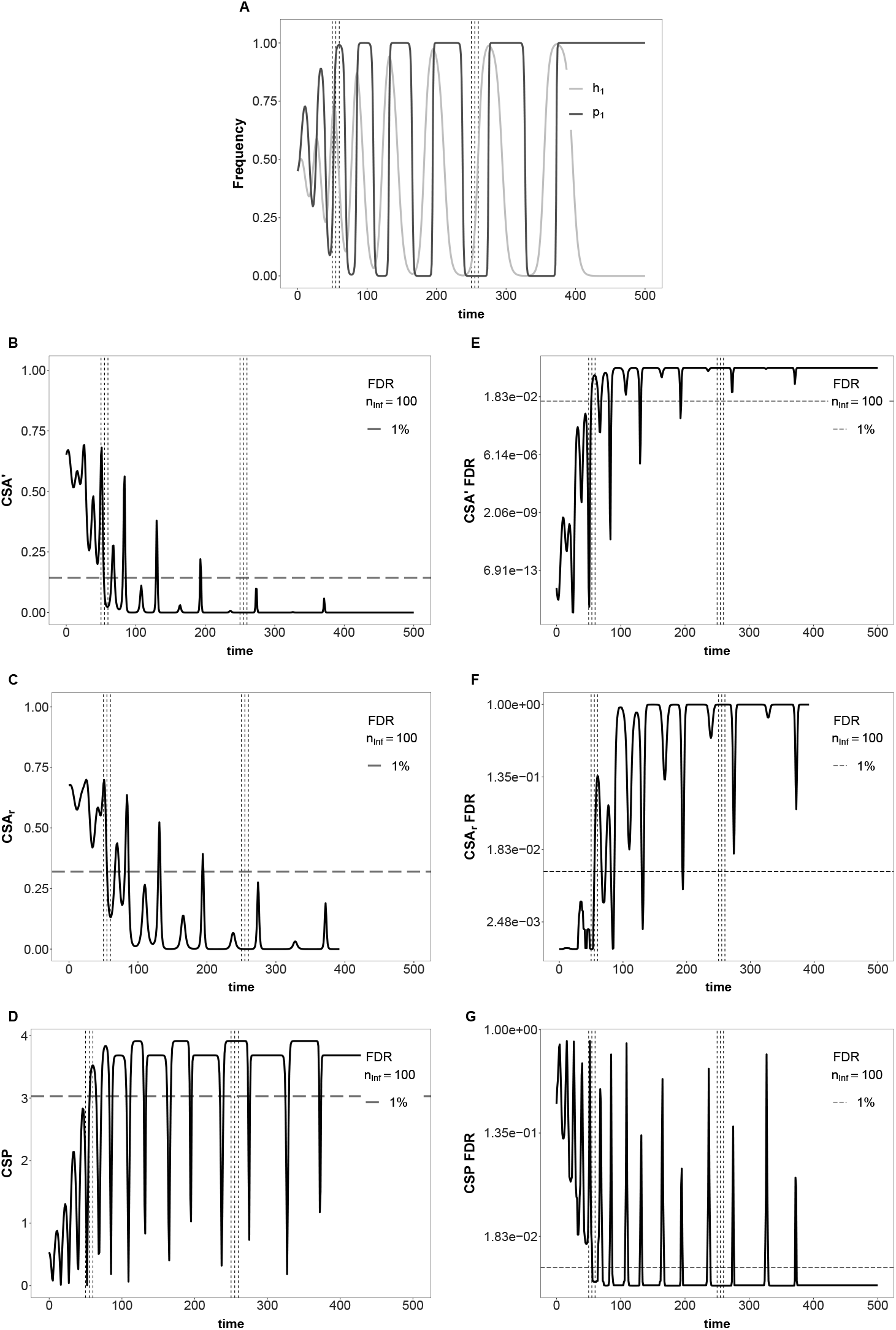
Temporal changes in allele frequencies, CSA′, CSA′_*r*_ and CSP in an unstable asymmetric MA-model (model A). Top: Temporal changes in allele frequencies. Left: Temporal changes of the three association indices. Right: corresponding FDR. Dashed lines correspond to a 1%-FDR, assuming *n_T_* = 200 and *n*_Inf_ = *n*_H_ = 100. The model parameters are *ω*_1_ = 0.9, *ω*_2_ = 0.7, *ϕ* = 0.8, *s* = 0.35, initial values *h*_1*,g*=0_ = *p*_1*,g*=0_ = 0.45. The y-axes in subfigures E, F and G have been log-transformed for better visualization of low FDR levels.

#### 3.2.2. Under a gene-for-gene infection matrix

For a gene-for-gene infection matrix (Table 1c) and for 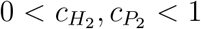 and 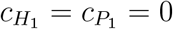 the non-trivial equilibrium frequencies are given by:

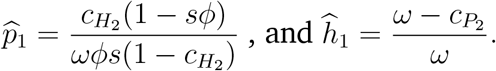

Inserting these frequencies into Eq. 10 and Eq. 11 we can obtain the values of the indices at the polymorphic equilibrium point.

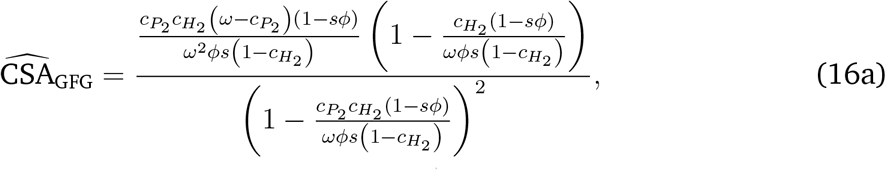

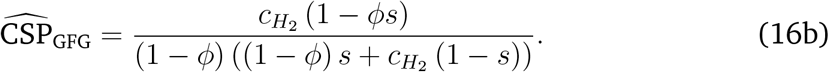

Note that under a gene-for-gene model (GFG), the equilibrium CSP values solely depend on host parameters 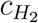, *ϕ* and *s* (independent of parasite cost 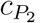), whereas the equilibrium CSA values depend on all the parameters in both species (Eq. 16, Fig. 3). The conditions under which CSP is defined are more restrictive under GFG than under MA. Unlike for MA, CSP is not defined when the exposed fraction is maximal (*ϕ* = 1) (see Eq. 11).

**Figure 3.**
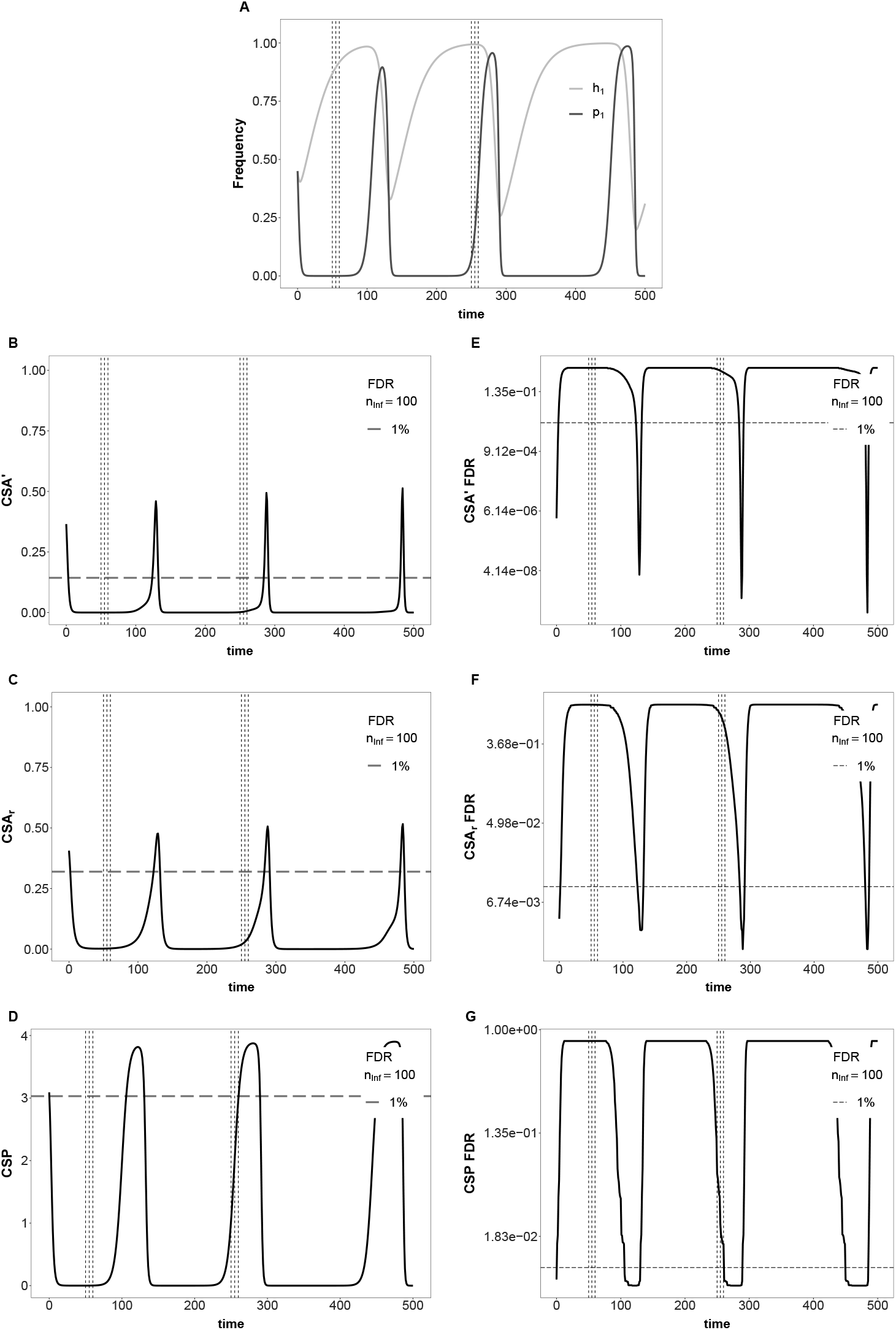
Temporal changes in allele frequencies, CSA′, CSA′_*r*_ and CSP in an unstable GFG-model (model A). Top: Temporal changes in allele frequencies. Left: Temporal changes of the three association indices. Right: corresponding FDR. Dashed lines correspond to a 1%-FDR assuming *n_T_* = 200 and *n*_Inf_ = *n*_H_ = 100. The model parameters are 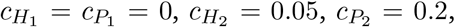 *ϕ* = 0.8, *s* = 0.35, *ω* = 0.9, initial values *h*_1*,g*=0_ = *p*_1*,g*=0_ = 0.45. The y-axes in subfigures E, F and G have been log-transformed for better visualization of low FDR levels.

### 3.3. Numerical simulations

#### 3.3.1. Arms race dynamics under model A

When simulating an asymmetric MA interaction for one representative parameter combination (*ω*_1_ = 0.9, *ω*_2_ = 0.7) over 500 generations (Fig. 2), coevolution results in arms race dynamics under Model A. We observe that CSA and CSP fluctuate over time due to the coevolutionary cycles and the corresponding allele frequency changes. Overall, CSA values decrease over time and result in high FDRs after *g* = 300 generations. Therefore, under unstable coevolutionary dynamics with increasing amplitude and period of the coevolutionary cycles (resulting ultimately in fixation of alleles), the associations between host and parasite alleles in the infected sample become too weak to be observable (Fig. 2b,c). Under an arms race dynamic, one host allele tends to reach a very high frequency and is subsequently tracked down by the parasite which generates the observed coevolutionary cycles. Along these cycles there is only a very limited amount of time at which both host and parasite alleles are found at intermediate frequencies yielding high values of CSA. On the other hand, the CSP values under the MA model are consistently high with very low FDR, exhibiting thus enough statistical power to detect the loci under coevolution (Fig. 2d,g). This demonstrates the importance of obtaining additional non-infected host samples. Under the same model A with a symmetric MA interaction, we find similar outcomes as in Fig. 2. The oscillations of CSA and CSP values perfectly match the allele frequency cycles and show a regular amplitude pattern (Fig. S8). Under the GFG model with arms race dynamics, CSA values are consistently small with narrow peaks (Fig. 3b,c) which are barely below the 1% FDR threshold (Fig. 3e,f). The CSP has several peaks at relatively low FDR (Fig. 3d,g). Yet, it maybe difficult to detect the coevolutionary loci even when time samples are available if a stringently low FDR is required (Fig. 3g). Comparing the results for a MA and a GFG-interaction under arms race dynamics shows that the combination of CSA and CSP values over time gives some indication about the (a)symmetry of the infection matrix, while the CSP exhibits the highest power to disentangle loci under coevolution from the neutral background. Generally, model A always results in arms race dynamics, however the amplitude and the time to fixation are affected by the underlying infection matrix and the coevolutionary costs.

#### 3.3.2. Trench warfare and arms race dynamics under model B

Under Model B, trench warfare dynamics can take place for MA-interactions and thus, allele frequencies can converge to a stable polymorphic equilibrium. Once the allele frequencies are at equilibrium, CSA accurately distinguishes the coevolving loci from the neutral background (Fig. 4b,c,e,f), while CSP decreases to zero (Fig. 4d). In Model B, the disease prevalence varies over time as a function of the changes in allele frequencies, so that CSP is expected to vary over time. However, once the allele frequencies have converged to a stable equilibrium the numbers in all host compartments eventually remain constant. Whenever the ratio of infected to non-infected individuals is the same for both host types the CSP drops to zero. The value of CSA at equilibrium depends on the respective equilib-rium frequencies and thus, is highest when alleles are at frequency 0.5 (Eq. 15) resulting in a a very low FDR. Even if coevolution results in stably sustained cycles, the CSA values would remain high as long as the amplitude of the cycles does not become too large and therefore alleles do not reach too high or too low frequencies (close to the boundaries). When allele frequency fluctuations do not occur and the system is already at the polymorphic equilibrium, CSA is fixed to a constant and high value (Fig. S9) with very low FDR, while the CSP is fixed to zero (under a symmetric MA infection matrix) with a maximum FDR. Note that the epidemiological model B can also generate arms race dynamics with a consequent fixation of one host and one parasite allele for some parameter combinations under a GFG-interaction (Fig. S10). In such cases, the obtained results are similar to those of Fig. 3 with both indices dropping to zero over time.

**Figure 4.**
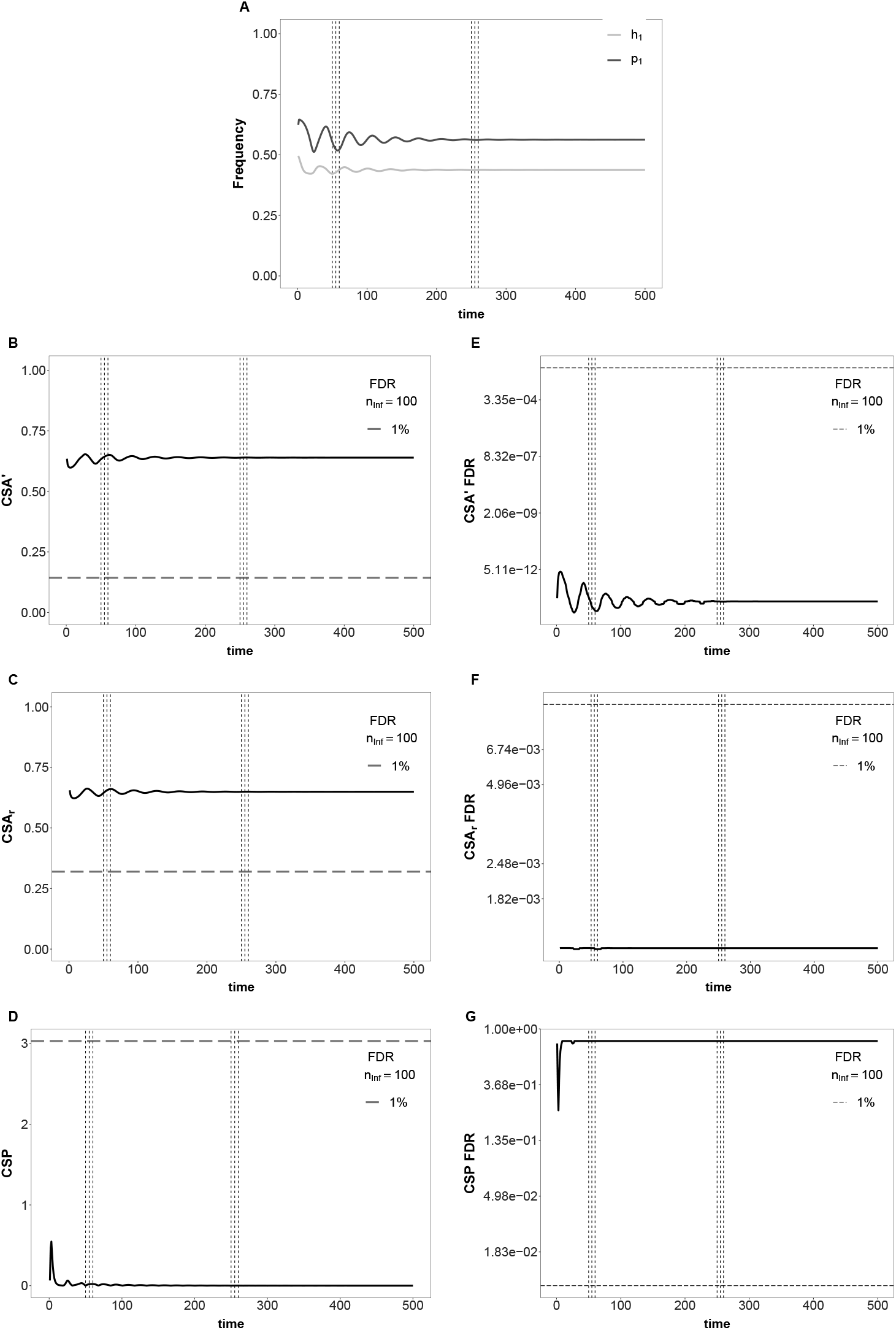
Temporal changes in allele frequencies, CSA′, CSA′_*r*_ and CSP in an epidemiological (model B) with an asymmetric MA-infection matrix. Top: Temporal changes in allele frequencies. Left: Tem-poral changes of the three association indices. Right: corresponding FDR. Dashed lines correspond to a 1%-FDR assuming *n_T_* = 200 and *n*_Inf_ = *n*_H_ = 100. The model parameters are *β* = 0.00005, *s* = 0.6, *ω*_1_ = 0.9, *ω*_2_ = 0.7, *b* = 1, *γ* = 0.9, and 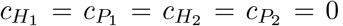. The initial values are *S*_1*,t*=0_ = *S*_2*,t*=0_ = 41500, *I*_11*,t*=0_ = *I*_12*,t*=0_ = *I*_21*,t*=0_ = *I*_22*,t*=0_ = 4150. The time interval for computations is *δ_t_* = 0.001. The y-axes in subfigures E, F and G have been log-transformed for better visualization of low FDR levels.

Generally, the parasite population is dying out when the disease transmission rate (*β*) is either too high or too low, irrespective of the underlying infection matrix. For intermediate disease transmission rates under a MA-interaction, we either observe stable limit cycles for large values of *s* (*s* ≈ 1) or convergence to a stable equilibrium for *s <* 1. These qualitative trends are similar when changing the infection matrix from MA to GFG, though the number of cycles to reach the stable equilibrium varies (Živković et al. 2019).

Therefore, the highlighted results from Model B reflect the most commonly observed dynamics in this model. Finally, we note that the two measures of CSA we introduce, CSA′ and CSA_*r*_ show the same trend and similar power under an arms race assumption, while the CSA_*r*_ is slightly more precise under trench warfare dynamics when allele frequencies reach very high or very low values (close to fixation or loss).

#### 3.3.3. Added value of time sampling

Our sampling schemes *Z*_1_ and *Z*_2_ fall into distinct parts of the allele frequency dynamics (Fig. 2-4 and Fig. S8-S10). For the analyzed asymmetric matching-alleles and symmetric matching-alleles interaction under model A the first interval *Z*_1_ is located around a peak with respect to CSA and CSP (Fig. 2 and Fig. S8) whereas it is located in a local mini-mum for the GFG interaction (Fig. 3). The second time interval *Z*_2_ coincides with very small values for the asymmetric MA-model (Fig. 2), the transition into a peak for the GFG model (Fig. 3) and a peak for the symmetric MA-model (Fig. S8). For the asymmetric MA-interaction under Model B, the first time interval encompasses a transition into a local maximum with respect to CSA with the allele frequencies still showing minor fluctuations but being already close to the equilibrium point (Fig. 4). Correspondingly, the CSP values are already very close to zero. The second time interval falls into the part of the dynamic where the allele frequencies have already converged to the equilibrium and hence CSA and CSP have also converged to constant values. The same pattern can be seen under a corresponding symmetric MA-interaction for both intervals *Z*_1_ and *Z*_2_ (Fig. S9). For the corresponding GFG-interaction both time intervals fall into the part of the dynamic with strong positive selection and therefore, both time intervals are characterized by low CSA and CSP values.

We also compare the *χ*^2^-tests and FDR values across time-samples within time intervals and between time intervals (Tab. 4-6 and Tab. S1-S3). Both, the symmetric and the asymmetric MA-allele interaction in Model B result in significant CSA-*χ*^2^-association tests irrespective of the time-point and interval considered, even if a stringent Bonferroni-correction is ap-plied (Tab. 6, Tab. S2). The corresponding FDRs for CSA′ are very low whereas the FDRs for CSA_*r*_ would result in large number of false positive when the number of comparisons is large. These results are in stark contrast to all interactions involving arms race dynamics (model A and a GFG-interaction for model B)(Tab. 4, Tab. 5, Tab. S1). For the asymmetric matching-alleles interaction in model A the CSA-*χ*^2^-test for *g* = 50 is significant where it is not for the other two time samples within the same interval *Z*_1_. This reflects the fact that the sample at *g* = 50 exactly falls on the peak for CSA values whereas the other two samples within the same interval already fall slightly outside the peak. On the other hand the CSP-*χ*^2^-tests for the other two time samples within *Z*_1_ are significant as are all three of them in *Z*_2_ (Tab. 4). The corresponding results for the symmetric MA are overall similar, though none of the time samples in the first interval is exactly located on the local maximum of the CSA-peak. The time samples for the GFG-dynamic under model A (Tab. 5, Fig. 3) and model B (Tab. S3) do not show any significant *χ*^2^-test for both intervals. We also note that in all studied examples the number of false positives, for both CSA and CSP, are fairly high if a large number of loci is to be analyzed irrespectively of the significance of the corresponding *χ*^2^-tests.

**Table 4.** Comparison of the *χ*^2^-test results and the FDRs corresponding to the values of CSA′, CSA_*r*_ and CSP for the six time samples taken under an asymmetric MA for model A (see Fig. 2). The model parameters are *ω*_1_ = 0.9, *ω*_2_ = 0.7, *ϕ* = 0.8, *s* = 0.35, with initial values *h*_1*,g*=0_ = *p*_1*,g*=0_ = 0.45 and sample sizes *n_T_* = 200 and *n*_Inf_ = 100. Asterisks (*) indicate significant p-values after Bonferroni correction for 10^7^ comparisons.

|  | g=50 | g=55 | g=60 | g=250 | g=255 | g=260 |
| --- | --- | --- | --- | --- | --- | --- |
| $\chi^2_{\text{CSA}}$ | 4.65e+01 | 1.11e+01 | 1.74e+00 | 6.41e-05 | 1.96e-06 | 6.71e-07 |
| p-value | 8.99e-12* | 8.82e-04 | 1.87e-01 | 9.94e-01 | 9.99e-01 | 9.99e-01 |
| $\text{CSA}'$ | 6.50e-01 | 7.80e-02 | 2.37e-02 | 9.34e-08 | 4.33e-09 | 3.53e-09 |
| FDR $\text{CSA}'$ | 6.38e-14 | 6.04e-02 | 3.51e-01 | 9.96e-01 | 9.96e-01 | 9.96e-01 |
| $\text{CSA}_r$ | 6.82e-01 | 3.33e-01 | 1.32e-01 | 8.00e-04 | 1.40e-04 | 8.19e-05 |
| FDR $\text{CSA}_r$ | 1.17e-03 | 9.51e-03 | 1.39e-01 | 9.96e-01 | 9.96e-01 | 9.96e-01 |
| $\chi^2_{\text{CSP}}$ | 3.46e+01 | 4.36e+01 | 5.83e+01 | 2.86e+01 | 7.35e+01 | 1.06e+02 |
| p-value | 3.98e-09 | 4.04e-11* | 2.27e-14* | 9.12e-08 | 1.03e-17* | 7.07e-25* |
| CSP | 1.75e+00 | 2.80e+00 | 3.52e+00 | 3.91e+00 | 3.91e+00 | 3.91e+00 |
| FDR CSP | 3.95e-02 | 1.58e-02 | 7.59e-03 | 7.04e-03 | 7.04e-03 | 7.04e-03 |

**Table 5.** Comparison of the *χ*^2^-test results and the FDRs corresponding to the values of CSA′, CSA_*r*_ and CSP for the six time samples taken under a GFG-interaction for model A (see Fig. 3). The model parameters are 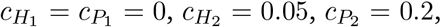, *ϕ* = 0.8, *s* = 0.35, *ω* = 0.9, with initial values *h*_1*,g*=0_ = *p*_1*,g*=0_ = 0.45, and sample sizes *n_T_* = 200 and *n*_Inf_ = 100.

|  | g=50 | g=55 | g=60 | g=250 | g=255 | g=260 |
| --- | --- | --- | --- | --- | --- | --- |
| $\chi^2_{\text{CSA}}$ | 3.74e-04 | 5.14e-04 | 7.94e-04 | 3.64e-02 | 8.73e-02 | 2.19e-01 |
| $p$-value | 9.85e-01 | 9.82e-01 | 9.78e-01 | 8.49e-01 | 7.68e-01 | 6.40e-01 |
| $\text{CSA}'$ | 1.47e-05 | 2.07e-05 | 3.27e-05 | 1.50e-03 | 3.21e-03 | 6.13e-03 |
| FDR $\text{CSA}'$ | 9.96e-01 | 9.96e-01 | 9.96e-01 | 8.91e-01 | 7.76e-01 | 6.80e-01 |
| $\text{CSA}_r$ | 1.93e-03 | 2.27e-03 | 2.82e-03 | 1.91e-02 | 2.95e-02 | 4.68e-02 |
| FDR $\text{CSA}_r$ | 9.93e-01 | 9.92e-01 | 9.90e-01 | 8.37e-01 | 7.14e-01 | 5.51e-01 |
| $\chi^2_{\text{CSP}}$ | 9.40e-08 | 2.28e-07 | 6.99e-07 | 2.13e-02 | 1.35e-01 | 7.58e-01 |
| $p$-value | 1.00e+00 | 1.00e+00 | 9.99e-01 | 8.84e-01 | 7.14e-01 | 3.84e-01 |
| CSP | 6.41e-04 | 1.10e-03 | 2.14e-03 | 1.04e+00 | 2.03e+00 | 2.98e+00 |
| FDR CSP | 8.04e-01 | 8.04e-01 | 8.04e-01 | 6.15e-02 | 2.33e-02 | 1.58e-02 |

**Table 6.** Comparison of the *χ*^2^-test results and the FDRs corresponding to the values of CSA′, CSA_*r*_ and CSP for the six time samples taken under an asymmetric MA for model B (see Fig. 4). The model parameters are *β* = 0.00005, *s* = 0.6, *ω*_1_ = 0.9, *ω*_2_ = 0.7, *b* = 1, *γ* = 0.9, and 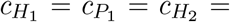 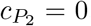. The initial values are *S*_1*,t*=0_ = *S*_2*,t*=0_ = 41500, *I*_11*,t*=0_ = *I*_12*,t*=0_ = *I*_21*,t*=0_ = *I*_22*,t*=0_ = 4150, the time interval for computations is *δ_t_* = 0.001, and sample sizes are *n_T_* = 200 and *n*_Inf_ = 100. Asterisks (*) indicate significant p-values after Bonferroni correction for 10^7^ comparisons.

|  | t=50 | t=55 | t=60 | t=250 | t=255 | t=260 |
| --- | --- | --- | --- | --- | --- | --- |
| $\chi^2_{\text{CSA}}$ | 4.16e+01 | 4.30e+01 | 4.36e+01 | 4.22e+01 | 4.22e+01 | 4.22e+01 |
| $p$-value | 1.15e-10* | 5.55e-11* | 3.96e-11* | 8.40e-11* | 8.19e-11* | 8.11e-11* |
| $\text{CSA}'$ | 6.33e-01 | 6.43e-01 | 6.50e-01 | 6.39e-01 | 6.40e-01 | 6.40e-01 |
| FDR $\text{CSA}'$ | 2.73e-13 | 1.02e-13 | 6.47e-14 | 1.74e-13 | 1.68e-13 | 1.68e-13 |
| $\text{CSA}_r$ | 6.45e-01 | 6.56e-01 | 6.61e-01 | 6.49e-01 | 6.50e-01 | 6.50e-01 |
| FDR $\text{CSA}_r$ | 1.18e-03 | 1.18e-03 | 1.17e-03 | 1.18e-03 | 1.18e-03 | 1.18e-03 |
| $\chi^2_{\text{CSP}}$ | 3.05e-06 | 6.34e-02 | 8.92e-02 | 1.30e-07 | 7.10e-05 | 8.74e-05 |
| $p$-value | 9.99e-01 | 8.01e-01 | 7.65e-01 | 1.00e+00 | 9.93e-01 | 9.93e-01 |
| CSP | 1.60e-04 | 1.78e-02 | 1.99e-02 | 2.76e-05 | 6.44e-04 | 7.15e-04 |
| FDR CSP | 8.04e-01 | 8.04e-01 | 8.04e-01 | 8.04e-01 | 8.04e-01 | 8.04e-01 |

## 4. Discussion

With technological advances, full genome sequencing of several hosts along with the parasite strains infecting them has become affordable. Recent studies (Ansari et al. 2017, Bartha et al. 2013, Lees et al. 2019) tested for the degree of association between different host and parasite SNPs by performing millions of pairwise comparisons to obtain a genome-wide natural co-GWAs. Here we develop two indices, CSA and CSP, measuring the strength of association between host and parasite SNPs. We demonstrate that the statistical power to locate the loci under coevolution depends on the 1) symmetry of the infection matrix, 2) type of dynamics, and 3) state of the allele frequency changes (whether a stable polymorphic equilibrium has been reached or not). We further show that CSA and CSP, provide different information with respect to these properties and can complement each other by having maximum power at different time points.

Based on previous studies it is expected that the power to detect coevolutionary loci varies in time due to the coevolutionary dynamics and the underlying allele frequency changes (Gandon and Nuismer 2009, MacPherson et al. 2018, Nuismer et al. 2017, Nuismer and Week 2019). Our approach is similar in spirit to studies measuring local adaptation by performing reciprocal transplant or common garden experiments across several host-parasite populations being connected by migration (Gandon and Nuismer 2009, Nuismer et al. 2017). The CSA measure is closely related to the covariance computed in Gandon and Nuismer (2009) and Nuismer et al. (2017), which also shows variable statistical power over time to detect coevolution. Therefore, we suggest obtaining samples from several time points, rather than a single time point, to extract as much information as possible. A similar idea was proposed in Gandon et al. (2008) for the study of local adaptation when using phenotypic (infection) data.

One crucial result of our study is the derivation of an indicative False Discovery Rate (FDR) for the association indices CSA and CSP. To do so, we assume that the analyzed samples originate from panmictic host and parasite populations with constant size. This allows us to apply population genetics theory to obtain the expected power and FDR of our indices for populations at drift-mutation equilibrium. In natural systems host and parasite population sizes (termed as the demographic history in population genetics) can change over time due to 1) eco-evolutionary feedback arising from the epidemiological dynamics (for example in model B), and/or due to 2) abiotic environmental factors such as resource availability and habitat suitability. It is possible to account for both type of demographic changes, by incorporating previous results on how they change the neutral SFS (Živković et al. 2019, 2015) into the calculation of the expected neutral distributions.

Further, it is known from a previous study (Živković et al. 2019) that the changes in neutral SFS under model B due to the coevolutionary cycles are very minor given that the initial population sizes are large enough. As we always consider large initial population sizes in our analysis, we therefore can roughly approximate the SFS by a stationary state SFS.

We assume that coevolution only involves one major bi-allelic locus per species. However, the MHC (major histocompatibility complex) loci in vertebrates (Horton et al. 2004) or NLRs (nucleotide-binding leucine rich repeat immune receptors) in plants (Van de Weyer et al. 2019) represent examples of gene families with complex genomic and functional structures which are likely to be involved into coevolution. We think that our indices are informative when several independent major loci (freely recombining and no epistatic or pleiotropic effects) are involved into the coevolutionary interaction as our indices are applied independently to each pair of host and parasite loci.

In the case of epistasis, it has been shown that positive epistatis increases the specificity of the parasite whereas negative epistasis widens the range of parasite infectivity (Ashby et al. 2014). Therefore, we expect the power of our indices to pinpoint the coevolving loci to increase (decrease) in the former case (in the latter case).

While we assume bi-allelic SNPs, several haplotypes can form the genetic basis of coevolution. Each haplotype can either consist of several alleles at different sites within a given gene (Seger 1988) or of bi-allelic SNPs across several linked loci (Tellier and Brown 2007a). In such cases the signature of coevolution and association (between host and parasite) would be weak for each single SNP and only visible once the defined haplotypes become the unit of analysis.

In principle, our indices can be extended to more than two-alleles per locus (Seger 1988) or to linked loci (Tellier and Brown 2007a), but theory suggests that, for these genetic structures, the coevolutionary dynamics become more complex (Seger 1988, Tellier and Brown 2007a).

We conclude by presenting a set of recommendations for applying this method to different host-parasite systems. One key requirement of this method is to keep track of the co-occurrence of host and parasite genotypes, which can be technically challenging especially in pool-sequencing approaches (as in Retel et al. 2019). It is advised to obtain random samples from infected and non-infected hosts and parasites at several time points from natural populations or from controlled coevolution experiments. Furthermore, sample sizes do affect the FDR and small sample sizes (*n_T_* < 25) decrease the power to detect the coevolving loci substantially (see Supplementary Online Material). Therefore, we suggest to collect large samples at few time points, rather than sampling small samples at several time points. It has been shown that polycyclic parasites, that is parasites with several infection cycles/generations per host generation, track down the host frequencies within a host generation (Tellier and Brown 2007b). Therefore, it is expected that the cross-species association method has a high power when applied to parasites with strong life-cycle dependency on their hosts. This effect should be strongly pronounced for parasites which have a much shorter generation times than their hosts, such as viruses (Ansari et al. 2017, Bartha et al. 2013) or bacterial pathogens of humans (Lees et al. 2019). For such types of parasites, taking serial samples within a single host generation should help to pinpoint coevolutionary loci very accurately. Here, the coevolving loci are expected to show an increasing association with the corresponding host loci over the course of a single host generation, especially if the infection matrix follows a MA model.

An implicit assumption in Ansari et al. (2017), Bartha et al. (2013), Bartoli and Roux (2017), Lees et al. (2019) and our model is that disease transmission is random and panmictic so that potentially every host can get into close contact with the disease (no population sub-structuring affecting disease transmission). Thus, before performing a cross-species association, it is crucial to assess and correct for 1) population sub-structure and kinship relationships within single populations, and 2) overall spatial structure. Additional bias can be caused by neglecting population structure when different populations are at different stages of the coevolutionary cycle (*e.g.* Gavrilets and Michalakis 2008, Sasaki 2000).

Co-infections are common for many diseases (Alizon et al. 2013, Tollenaere et al. 2016). A solution to deal with such multi-strain infections in a co-GWAS-setting is to base the analysis on the major parasite strain found on the infected host (Bartha et al. 2013). The independence between neutral and coevolving loci can be violated for viruses, bacteria or clonal fungi due to the potential absence of recombination. This problem can addressed by following the approach outlined in Bartha et al. (2013), by first inferring a phylogeny of the virus samples, followed by a subsequent identification of clusters of polymorphic SNPs across the phylogeny and performing the co-GWAs based on the identified clusters (Ansari et al. 2017, Bartha et al. 2013).

Host-parasite coevolution is a multifaceted process. We have shown that natural co-GWAs have the power to pinpoint genes involved in this process especially when time-samples are available, while we acknowledge potential shortcomings due to model assumptions, and dependency on the underlying infection matrix and coevolutionary allele frequency changes. The approach presented here is also potentially applicable to detect major genes which are determining the compatibility between symbionts in mutualistic interactions. A further worthwhile development will be the explicit inclusion of spatial structure of host and parasite populations.

## Acknowledgments

This work was funded by the German Research Foundation (DFG). HM received support from grant numbers TE809/3-1 and 4-1 (awarded to Aurélien Tellier) within the DFG Priority Programme 1819. SJ received support from grant number TE809/3-2 (awarded to Aurélien Tellier) within the DFG Priority Programme 1819 and grant number TE809/6-2 (awarded to Aurélien Tellier and Wolfgang Stephan) within the DFG Priority Programme 1590.

## Data accessibility statement

A GitHub repository is being created with all the codes used for simulating data, calculating distributions and analyzing results. This will be made publicly available upon acceptance.

## Author contributions

All three authors (HM, AT and SJ) designed the study, implemented the simulations, analyzed the results, wrote the manuscript and revised the submitted version of the manuscript.

