## Supplementary material for "Cross-Species association statistics for genome-wide studies of host and parasite polymorphism data": Online Supplement

Short title: Host-parasite associations statistics

Hanna Märkle[a,b,1], Aurélien Tellier[a,1,*], Sona John[a,1,*]

[a]*Professorship for Population Genetics, Department of Life Science Systems, Technical University of Munich, 85354 Freising, Germany*
[b]*now Department of Ecology and Evolution, University of Chicago, 1101 E 57th St, Chicago, IL 60637, USA*

[1]All three authors contributed equally.

### S1. The neutral site frequency spectrum

We assume for simplicity that the allele frequency distribution of neutral SNPs in both, the host and the parasite, follows the site frequency spectrum (SFS) under drift-mutation equilibrium for a Wright-Fisher model under constant population size. Further, we assume that there is no outgroup sequence available, thus the ancestral and derived state are unknown for a given SNP. The expected folded SFS $\eta = \{\eta_1, \ldots, \eta_{\lfloor n/2 \rfloor}\}$ under drift-mutation equilibrium for a sample of size $n$ is given by (Durrett 2010, p50):

$$\eta_k = \frac{\frac{\theta}{k} + \frac{\theta}{n-k}}{1 + \delta_{k,n-k}} \quad \text{for } 1 \leq k \leq \lfloor n/2 \rfloor \tag{S1}$$

where $\theta$ is the population mutation rate, $\lfloor n/2 \rfloor$ denotes the largest integer being smaller or equal to $n/2$ and $\delta_{k,l}$ is Kronecker's delta with

$$\delta_{k,l} = \begin{cases} 0 & \text{for } k \neq l \\ 1 & \text{for } k = l. \end{cases}$$

Thus, the probability ($p_k$) to choose a SNP with minor allele frequency $k$ in a sample of size $n$, is given by:

$$p_k = \frac{\eta_k}{\sum_{i=1}^{\lfloor n/2 \rfloor} \eta_i} = \frac{\left( \frac{\frac{1}{k} + \frac{1}{n-k}}{1 + \delta_{k,n-k}} \right)}{\sum_{i=1}^{\lfloor n/2 \rfloor} \left( \frac{\frac{1}{k} + \frac{1}{n-k}}{1 + \delta_{k,n-k}} \right)}. \tag{S2}$$

These probabilities are independent of the population size and the mutation rate.

Our computations include singletons, that is alleles with frequency $1/n$ in the sample. However, it is known that the sequencing and detection of singletons can be biased (*e.g.* with NGS technologies or pooling of samples). Therefore, singletons can be also removed from the CSA calculation and the CSP calculations should be adjusted accordingly by constraining the minor allele frequency in the infected subsample to be at least equal to two. Furthermore, if more complex demographic scenarios want to be included via their influence on the neutral host and/or parasite SFS, the probability S2 can be adjusted analytically or from results of coalescent simulations (Živković et al. 2019, 2015).

### S2. CSA and CSP calculations

In this supplement we add some details on the computations of the CSA and CSP values for neutral loci as well as additional figures.

#### S2.1. Cross species association index (CSA)

As we describe in the main text, we have obtained $n_{\text{Inf}}$ host samples and one representative parasite strain from each of these infected hosts. Thus, the host sample size ($n_{\text{Inf}}$) and the parasite sample size ($n_{\text{Par}}$) are the same ($n = n_{\text{Inf}} = n_{\text{par}}$). In order to compute the expected CSA for neutral SNPs we first have to derive an expression for the expected value of CSA ($E(\text{CSA}_{vw})$) measuring the association between a host SNP with minor allele frequency $v$ and a parasite SNP with minor allele frequency $w$. Therefore, we first compute the number of all such possible combinations. For each combination, the value CSA is $\text{CSA}_{vw,k}$ and the probability of that particular combination is $\binom{v}{k}\binom{n_{\text{Inf}}-v}{w-k}$. The expectation $E(\text{CSA}_{vw})$ is then:

$$\begin{aligned} E(CSA_{vw}) &= \Omega_{vw} \sum_{k=0}^{l} \frac{\binom{v}{k}\binom{n_{\text{Inf}}-v}{w-k}}{\binom{n_{\text{Inf}}}{w}} CSA_{vw,k}, & \text{(S3)} \\ &= \Omega_{vw} \sum_{k=0}^{l} \frac{\binom{v}{k}\binom{n_{\text{Inf}}-v}{w-k}}{\binom{n_{\text{Inf}}}{w}} \left( \left| \frac{k}{n_{\text{Inf}}} \cdot \frac{n_{\text{Inf}} - v - (w-k)}{n_{\text{Inf}}} - \frac{v-k}{n_{\text{Inf}}} \cdot \frac{w-k}{n_{\text{Inf}}} \right| \right), & \text{(S4)} \\ &= \Omega_{vw} \sum_{k=0}^{l} \frac{\binom{v}{k}\binom{n-v}{w-k}}{\binom{n_{\text{Inf}}}{w}} \left| \frac{k n_{\text{Inf}} - vw}{n_{\text{Inf}}^2} \right|, & \text{(S5)} \end{aligned}$$

where $l = min(v, w)$.

Here the index $k$ can be interpreted as the number of hosts with the minor allele which are infected by a parasite with the minor allele, or put in a different way, $w - k$ out of the $n_{\text{Inf}} - v$ hosts with the major allele are infected by a parasite with the minor allele.

Accordingly, $v-k$ hosts with the minor allele are infected by a parasite which has the major allele, and $n_{\text{Inf}} - v - (w-k)$ hosts with the major allele are infected by parasites with the major allele. We define $\Omega_{vw}$ as the normalization for either obtaining CSA$'$ (with $\Omega_{vw} = 4$, as in Eq. 2) or CSA$_r$ (with ($\Omega_{vw} = \frac{1}{\sqrt{\frac{v}{n_{\text{Inf}}}\frac{n-v}{n_{\text{Inf}}}\frac{w}{n_{\text{Inf}}}\frac{n-w}{n_{\text{Inf}}}}}$), from Eq. 3).

#### S2.2. Distribution of CSA for different sample sizes

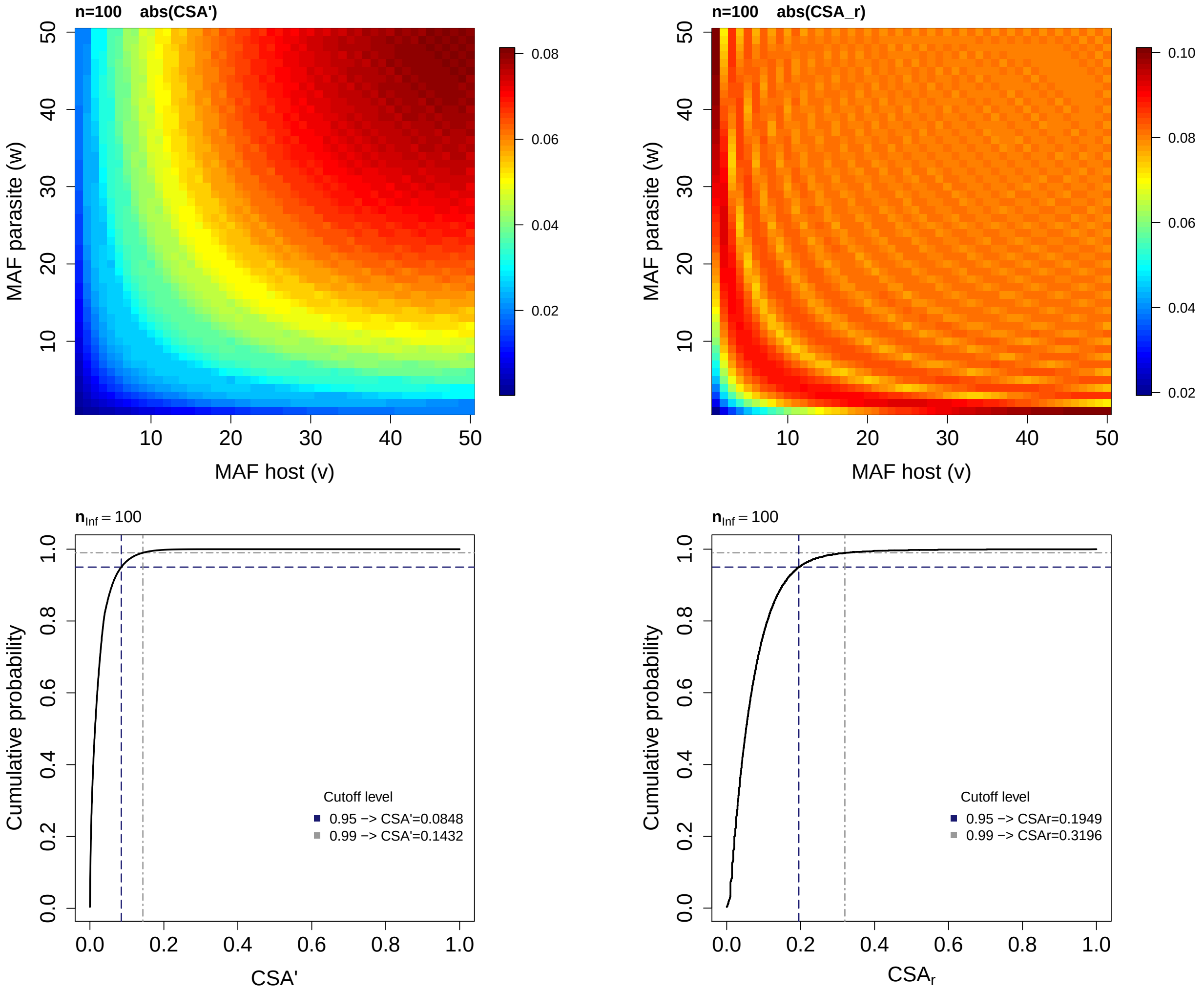


Figure S1 Expected values of CSA′ (top left) and $\text{CSA}_r$ (top right) when comparing all neutral host SNPs with minor allele frequency $v$ ($v \in \{1, \ldots, \lfloor n_{\text{Inf}}/2 \rfloor\}$) to all neutral parasite SNPs with minor allele frequency $w$ ($w \in \{1, \ldots, \lfloor n_{\text{Par}}/2 \rfloor\}$ and the resulting expected cumulative distribution function of E(CSA') (bottom left) and $\text{E}(\text{CSA}_r)$ (bottom right) for a sample size of $n_{\text{Inf}} = n_{\text{Par}} = 100$.

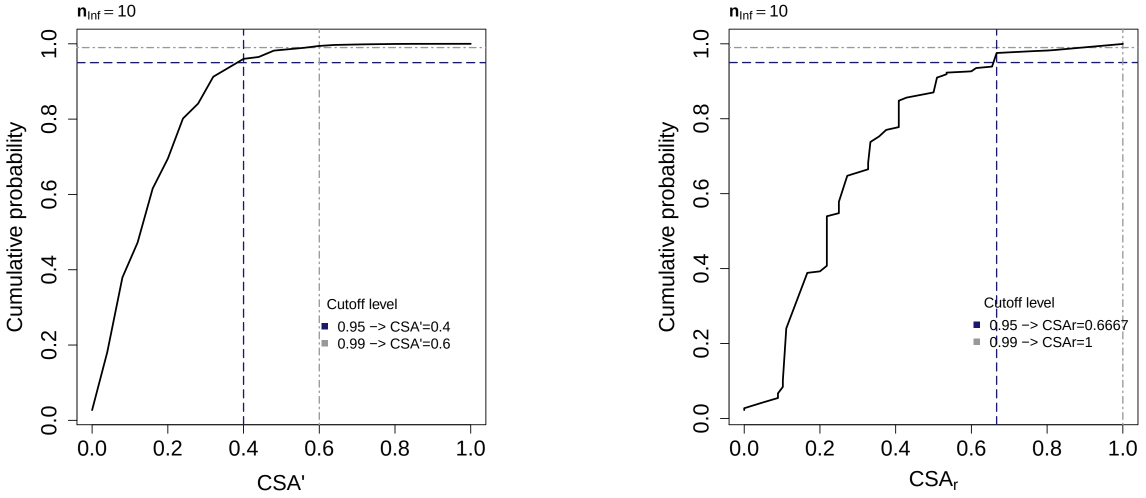


Figure S2  Expected cumulative distribution function of CSA′ (left) and $\text{CSA}_r$ (right) when comparing all neutral host SNPs with minor allele frequency $v$ ($v \in \{1, \ldots, \lfloor n/2 \rfloor\}$) to all neutral parasite SNPs with minor allele frequency $w$ ($w \in \{1, \ldots, \lfloor n/2 \rfloor\}$ for a sample size of $n_{\text{Inf}} = n_{\text{Par}} = 10$.

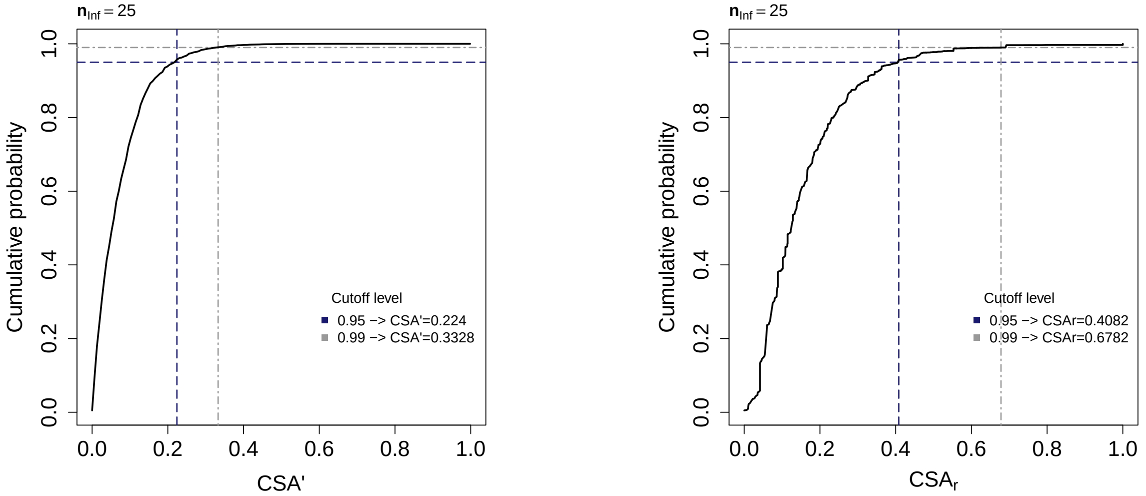


Figure S3  Expected cumulative distribution function of CSA′ (left) and $\text{CSA}_r$ (right) when comparing all neutral host SNPs with minor allele frequency $v$ ($v \in \{1, \ldots, \lfloor n/2 \rfloor\}$) to all neutral parasite SNPs with minor allele frequency $w$ ($w \in \{1, \ldots, \lfloor n/2 \rfloor\}$ for a sample size of $n_{\text{Inf}} = n_{\text{Par}} = 25$.

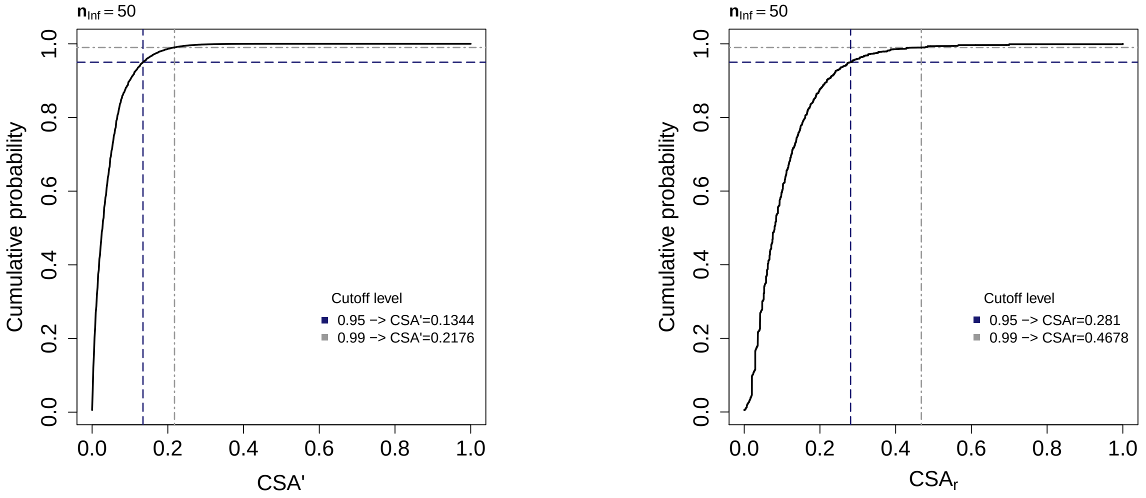


Figure S4 Expected cumulative distribution function of CSA$'$ (left) and CSA$_r$ (right) when comparing all neutral host SNPs with minor allele frequency $v$ ($v \in \{1, \ldots, \lfloor n/2 \rfloor\}$) to all neutral parasite SNPs with minor allele frequency $w$ ($w \in \{1, \ldots, \lfloor n/2 \rfloor\}$ for a sample size of $n_{\text{Inf}} = n_{\text{Par}} = 50$.

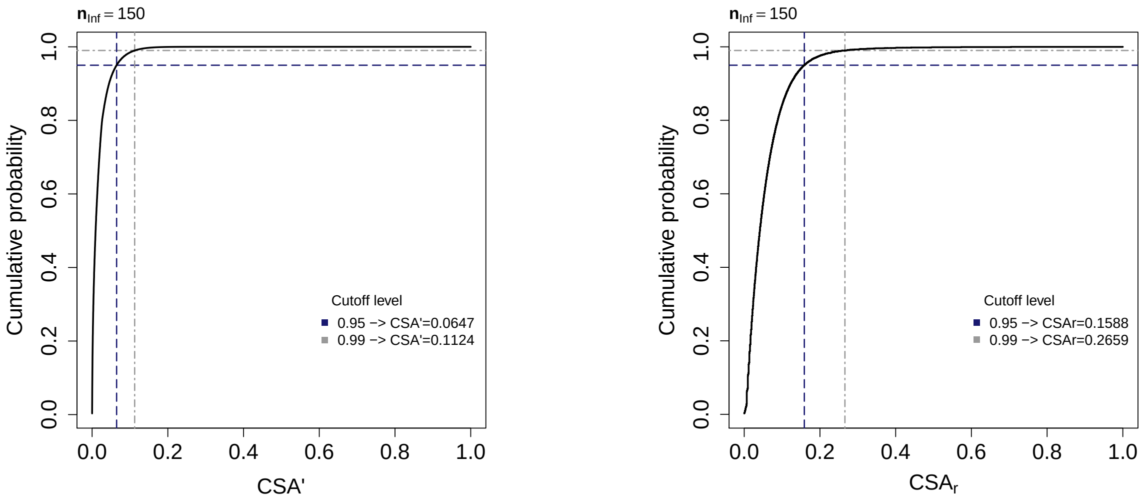


Figure S5 Expected cumulative distribution function of CSA$'$ (left) and CSA$_r$ (right) when comparing all neutral host SNPs with minor allele frequency $v$ ($v \in \{1, \ldots, \lfloor n/2 \rfloor\}$) to all neutral parasite SNPs with minor allele frequency $w$ ($w \in \{1, \ldots, \lfloor n/2 \rfloor\}$ for a sample size of $n_{\text{Inf}} = n_{\text{Par}} = 150$.

Note that the $0.95$ and $0.99$ cut-offs levels obtained from the cumulative distribution function for the CSA values translate directly into $5\%$ and $1\%$ false discovery rates (FDR). Similarly, an FDR value is computed from the neutral distribution for each CSA value at coevolving loci and represented in the main text and SI figures.

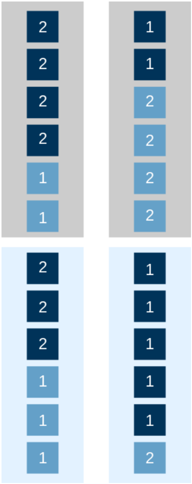


Figure S6  Two possible host configurations when sampling a total number $n_{\mathrm{T}} = 12$ host individuals among which $n_{\mathrm{Inf}} = 6$ individuals are infected (grey box) and $n_{\mathrm{H}} = 6$ individuals are healthy (light blue box) and the minor host allele frequency is $v = 5$. Host individuals which have the minor allele (based on the whole sample) are shown in light blue, host individuals with the major allele (based on the whole sample) are shown in dark blue. Labelling of the alleles for the calculation of CSP is based on the minor allele frequency in the infected subsample. On the left, the minor allele is labelled by $1$ as it is also the minor allele in the infected subsample. On the right, the major allele of the total sample is labelled by $1$ as it represents the allele with minor allele frequency in the infected subsample.

#### S2.3. Cross species prevalence index (CSP)

We label the host allele with minor frequency in the **infected subsample** as $i = 1$ and the host allele with major frequency in the infected subsample as $i = 2$. Note that the allele with minor allele frequency in the infected subsample is not necessarily the minor allele in the whole sample (see Fig. S6). In cases where both alleles have equal frequencies in the infected subsample, the allele with minor allele frequency in the whole sample will be labelled as $1$ and the allele with major allele frequency in the whole sample will be labelled as $2$. Therefore, $F_{11}$ ($F_{12}$) is the proportion of hosts with label $1$ which are infected by a parasite with the minor (major) allele frequency. $F_{21}$ ($F_{22}$) is the proportion of hosts with label $2$ which are infected by a parasite with the minor (major) allele frequency. Further, $F_{10}$ (respectively $F_{20}$) is the proportion of non-infected hosts carrying allele $1$ (respectively $2$). If for a neutral locus there are $v$ minor alleles in the total host sample $n_T$, these minor alleles can be found with equal probability on each of the $n_{\mathrm{T}}$ individuals (irrespective of the infection status) as a neutral SNP does not have an effect on the infection outcome. Similarly, all of the $w$ parasite minor alleles can be randomly assigned to any of the $n_{\text{par}}$ parasite individuals which are infecting the $n_{\text{Inf}}$ host individuals. Note that CSP is only informative when the minor and major allele can be found in both, the infected and the non-infected subsample. Therefore, we exclude SNPs which are singletons in the total host sample ($n_{\text{T}}$). We proceed as follows to obtain the expected CSP for a neutral host SNP with minor allele frequency $v$ and and neutral parasite SNP with minor allele frequency $w$.

First, we have to find all host combinations (and their probability) for which the minor and major host alleles are found in both the infected and non-infected subsamples. We define $z$ as the number of minor host alleles which are found in the infected subsample for a given combination. Accordingly, the number of major host alleles in the infected subsample is $n_{\text{Inf}} - z$, the number of minor host alleles in the non-infected subsample is $v - z$ and the number of major host alleles in the non-infected subsample is $n_T - n_{\text{Inf}} - (v - z)$. Based on the resulting composition of the infected subsample the alleles are labelled. The indicator variable $\lambda$ is used to keep track of whether the minor allele in the total sample is the minor ($\lambda = 0$) or the major ($\lambda = 1$) allele in the infected subsample.

Second, the $n_{\text{par}}$ parasites among which $w$ individuals have the minor parasite allele are assigned within the $n_{\text{Inf}}$ sample. Hereby, $k$ denotes the number of hosts with label $1$ which are infected by a parasite with the minor allele (see CSA). Thus, the expected value of CSP for a SNP with minor allele frequency $v$ in the host and minor allele frequency $w$ in the parasite is given by:

$$E(CSP_{vw}) = \sum_{z=\rho}^{m-1} \frac{\binom{n_{\text{Inf}}}{z}\binom{n_H}{v-z}}{\binom{n_{\text{T}}}{v} - \sum\limits_{b=0}^{\rho-1}\binom{n_{\text{Inf}}}{b}\binom{n_H}{v-b} - \sum\limits_{b=m}^{v}\binom{n_{\text{Inf}}}{b}\binom{n_H}{v-b}} \sum_{k=0}^{min(w,a)} \frac{\binom{a}{k}\binom{n_{\text{Inf}}-a}{w-k}}{\binom{n_{\text{Inf}}}{w}} \left| \frac{\frac{k}{n_{\text{T}}} + \frac{a-k}{n_{\text{T}}}}{\frac{\lambda n_H + (v-z)(-1)^{\lambda}}{n_{\text{T}}}} - \frac{\frac{w-k}{n_{\text{T}}} + \frac{n_{\text{Inf}}-a-(w-k)}{n_{\text{T}}}}{\frac{(1-\lambda)n_H - (v-z)(-1)^{\lambda}}{n_{\text{T}}}} \right| \tag{S6}$$

$$E(CSP_{vw}) = \sum_{z=\rho}^{m-1} \frac{\binom{n_{\text{Inf}}}{z}\binom{n_H}{v-z}}{\binom{n_{\text{T}}}{v} - \sum\limits_{b=0}^{\rho-1}\binom{n_{\text{Inf}}}{b}\binom{n_H}{v-b} - \sum\limits_{b=m}^{v}\binom{n_{\text{Inf}}}{b}\binom{n_H}{v-b}} \sum_{k=0}^{min(w,a)} \frac{\binom{a}{k}\binom{n_{\text{Inf}}-a}{w-k}}{\binom{n_{\text{Inf}}}{w}} \left| \frac{a}{\lambda n_H + (v-z)(-1)^{\lambda}} - \frac{n_{\text{Inf}} - a}{(1-\lambda)n_H - (v-z)(-1)^{\lambda}} \right|$$

$$E(CSP_{vw}) = \sum_{z=\rho}^{m-1} \frac{\binom{n_{\text{Inf}}}{z}\binom{n_H}{v-z}}{\binom{n_{\text{T}}}{v} - \sum\limits_{b=0}^{\rho-1}\binom{n_{\text{Inf}}}{b}\binom{n_H}{v-b} - \sum\limits_{b=m}^{v}\binom{n_{\text{Inf}}}{b}\binom{n_H}{v-b}} \left| \frac{a}{\lambda n_H + (v-z)(-1)^{\lambda}} - \frac{n_{\text{Inf}} - a}{(1-\lambda)n_H - (v-z)(-1)^{\lambda}} \right| \tag{S7}$$

$\rho = max(1, v - n_H + 1)$, $m = min(n_{\text{Inf}}, v)$, $a = min(z, n_{\text{Inf}} - z)$

$$\lambda = \begin{cases} 0 & \text{for } z \leq n_{\text{Inf}} - z \\ 1 & \text{for } z > n_{\text{Inf}} - z \end{cases} \tag{S8}$$

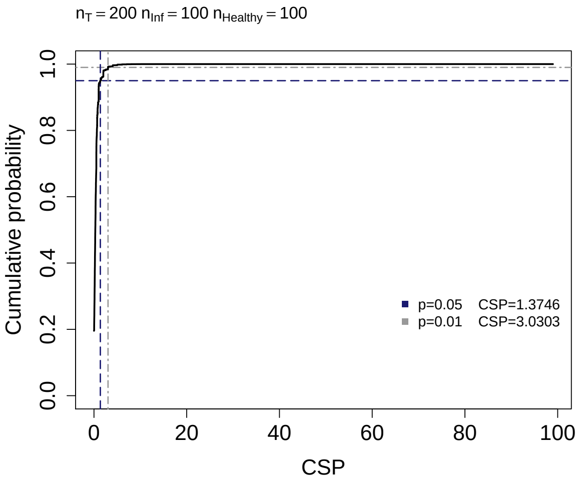


Figure S7 Cumulative distribution function of the expected value of CSP when taking a host sample of total size $n_{\mathrm{T}} = 200$ which includes $n_{\mathrm{Inf}} = 100$ infected hosts and $n_{\mathrm{H}} = 100$ healthy hosts.

The condition that $z$ starts from $\rho = max(1, v - (n_T - n_{\mathrm{Inf}}) + 1)$ is necessary to avoid combinations where 1) no minor allele is found in the infected subsample, and 2) the healthy sample only consists of hosts with the minor allele. The condition that $z$ has values up to $m - 1$ is necessary to avoid two configurations where 1) none of the minor alleles is found in the non-infected subsample, and 2) all individuals in the infected subsample have the minor allele. For a given host allele combination, we perform the labelling step mentioned above by defining $a = min(z, n_{\mathrm{Inf}} - z)$. Then, we assign the $n_{\mathrm{Par}}$ parasites, $w$ of them having the minor allele, to the $n_{\mathrm{Inf}}$ host. Here, $k$ is the number of hosts with label $1$ which are infected by a parasite with the minor allele $(F_{11} \cdot n_T)$. Accordingly, $a - k$ is the number of hosts with label $1$ which are infected by a parasite with major allele $(F_{12} \cdot n_T)$, $w - k$ is the number of hosts with label $2$ which are infected by a parasite with minor allele $(F_{21} \cdot n_T)$ and $n_{\mathrm{Inf}} - a - (w - k)$ with label $2$ which are infected by a parasite with the major allele. Note that the $0.95$ and $0.99$ cut-offs levels obtained from the cumulative distribution function for the CSP values (see Fig. S7) translate directly into $5\%$ and $1\%$ false discovery rates (FDR). Similarly, an FDR value is computed from the neutral distribution for each CSP value at coevolving loci and represented in the main text and SI figures.

### S3. Supplementary figures

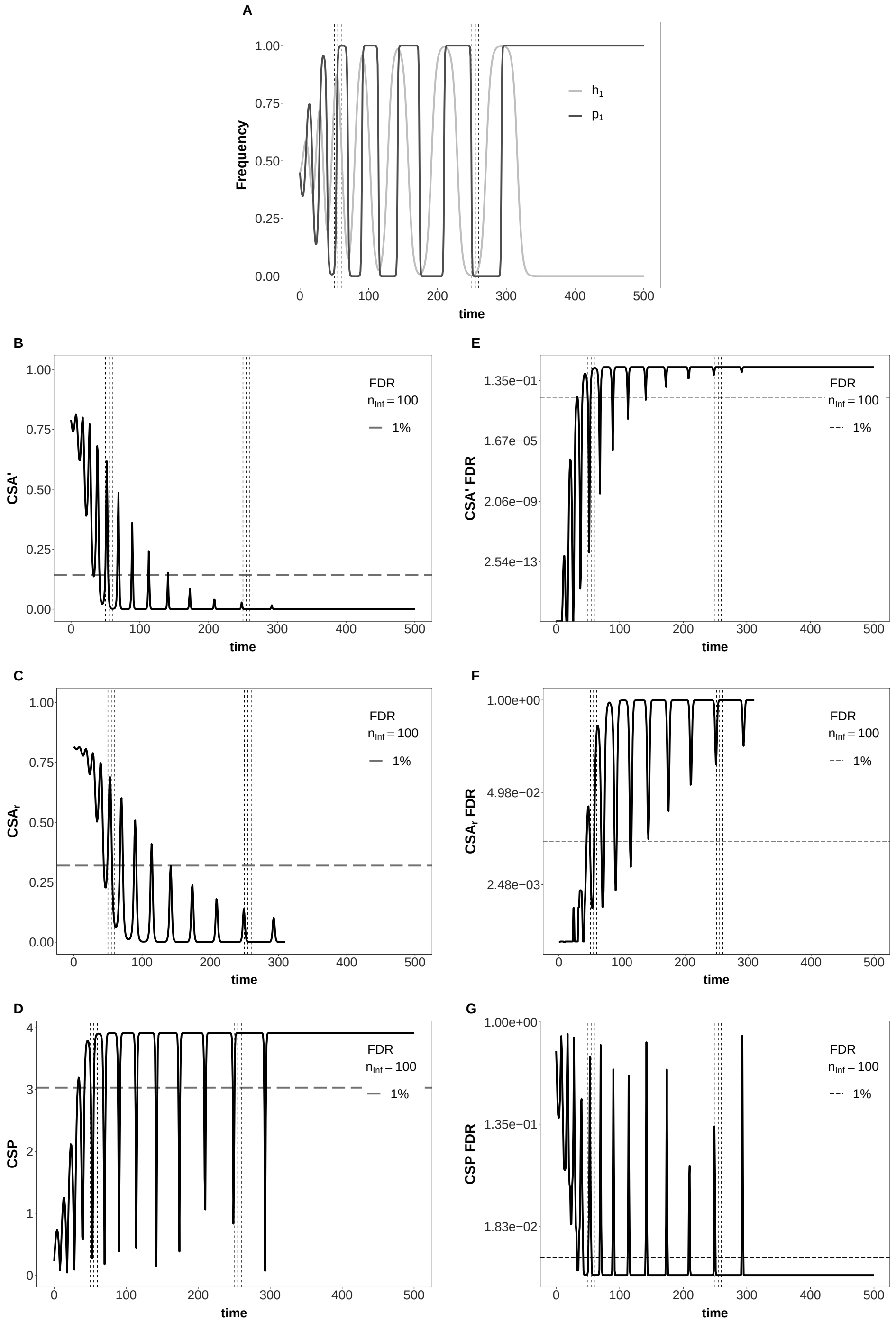


Figure S8 Temporal changes in allele frequencies, $\text{CSA}', \text{CSA}'_r$ and CSP in an unstable MA-model (model A) with one parasite generation per host generation. Top: Temporal changes in allele frequencies. Left: Temporal changes of the three association indices. Right: corresponding FDR. Dashed lines correspond to a $1\%$-FDR, assuming $n_T = 200$ and $n_{\text{Inf}} = n_{\text{H}} = 100$. The parameters values of the model are: $\omega_1 = \omega_2 = 0.9$, $c_{H_1} = c_{P_1} = c_{H_2} = c_{P_2} = 0$, $\phi = 0.8$, $s = 0.35$, $h_{1,\text{init}} = p_{1,\text{init}} = 0.45$. The y-axes in subfigures E, F and G have been log-transformed for better visualization of low FDR levels.

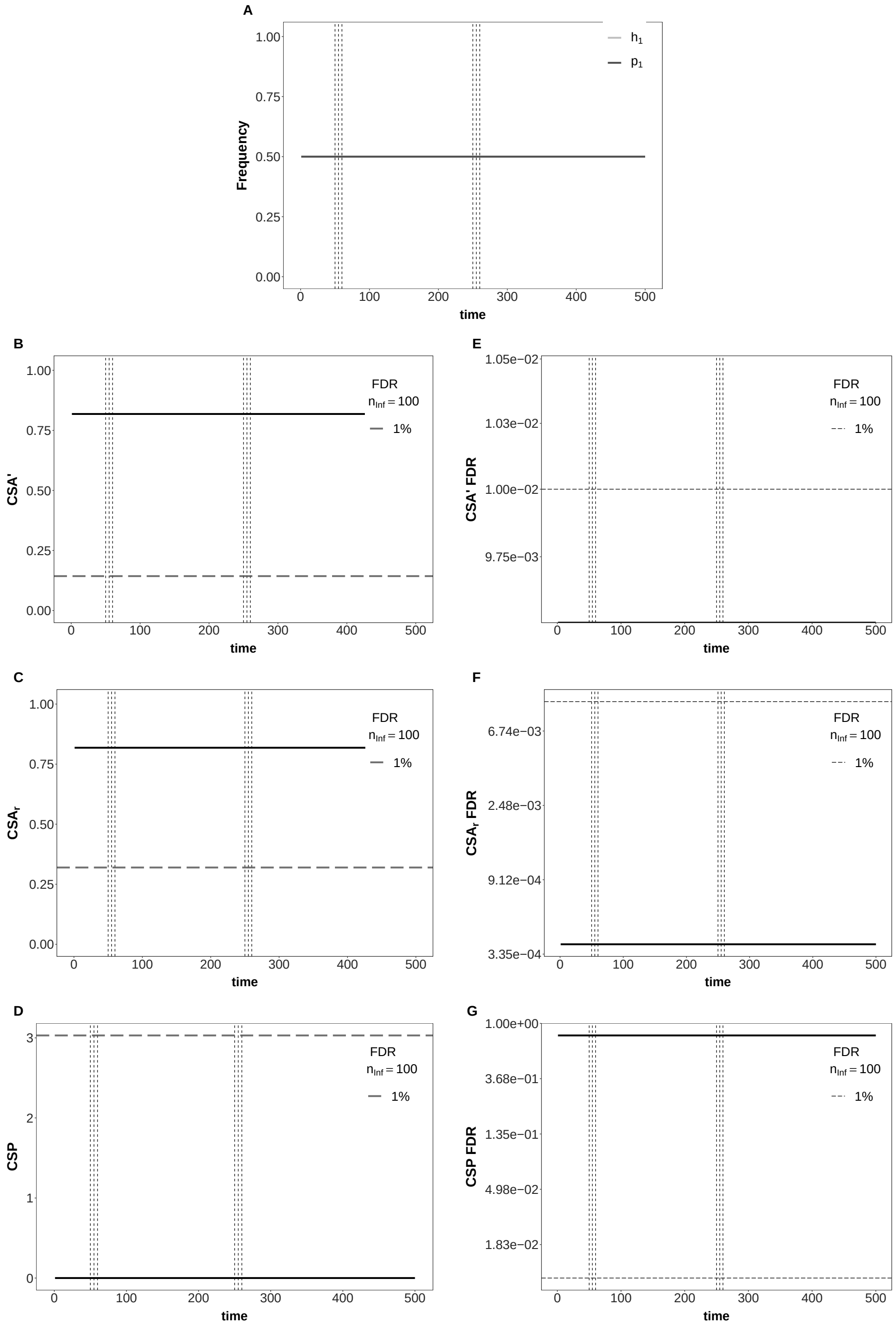


Figure S9 Temporal changes in allele frequencies, CSA$'$, CSA$'_r$ and CSP in an epidemiological (model B) with a symmetric MA-infection matrix. Top: Temporal changes in allele frequencies. Left: Temporal changes of the three association indices. Right: corresponding FDR. Dashed lines correspond to a 1%-FDR, assuming $n_T = 200$ and $n_{\text{Inf}} = n_{\text{H}} = 100$. The parameters values of the model are: $c_{H_1} = c_{P_1} = c_{H_2} = c_{P_2} = 0$, $\beta = 0.00005$, $s = 0.6$, $\omega_1 = \omega_2 = 0.9$, $S_{1,\text{init}} = S_{2,\text{init}} = 41500$, $I_{11} = I_{12} = I_{21} = I_{22} = 4150$, $\delta_t = 0.001$, $b = 1$, $\gamma = 0.9$. The y-axes in subfigures E, F and G have been log-transformed for better visualization of low FDR levels.

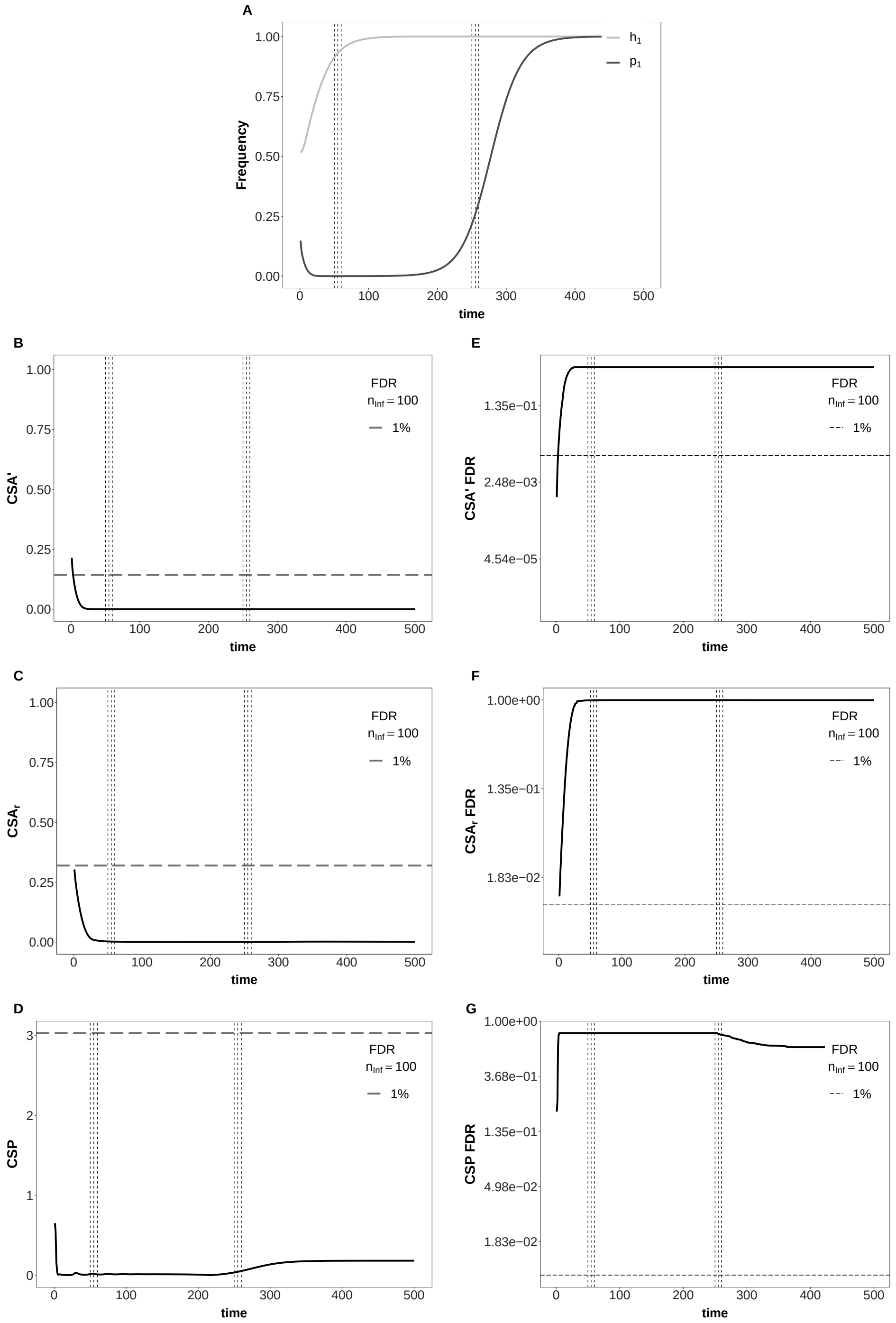


Figure S10 Temporal changes in allele frequencies, CSA$'$, CSA$'_r$ and CSP in an epidemiological (model B) with an GFG-infection matrix. Top: Temporal changes in allele frequencies. Left: Temporal changes of the three association indices. Right: corresponding FDR. Dashed lines correspond to a 1%-FDR, assuming $n_T = 200$ and $n_{\text{Inf}} = n_{\text{H}} = 100$. The parameters values of the model are: $c_{H_1} = c_{P_1} = 0$, $c_{H_2} = c_{P_2} = 0.05$, $\beta = 0.00005$, $s = 0.6$,$\omega = 0.9$, $S_{1,\text{init}} = S_{2,\text{init}} = 41500$, $I_{11} = I_{12} = I_{21} = I_{22} = 4150$, $\delta_t = 0.001$, $b = 1$, $\gamma = 0.9$. The y-axes in subfigures E, F and G have been log-transformed for better visualization of low FDR levels.

### S4. Supplementary tables

Table S1 Comparison of the $\chi^2$-test results and the FDRs corresponding to the values of CSA', CSA$_r$ and CSP for the six time samples taken under an symmetric MA for model A (see Fig. S8). The model parameters are $\omega_1 = 0.9$, $\omega_2 = 0.9$, $\phi = 0.8$, $s = 0.35$, with initial values $h_{1,g=0} = p_{1,g=0} = 0.45$, and sample sizes $n_T = 200$ and $n_{\text{Inf}} = 100$. Asterisks (*) indicate significant p-values after Bonferroni correction for $10^7$ comparisons.

| | g=50 | g=55 | g=60 | g=250 | g=255 | g=260 |
|---|---|---|---|---|---|---|
| $\chi^2_{\text{CSA}}$ | 1.58e+01 | 2.57e+01 | 5.21e-01 | 1.45e+00 | 2.08e-04 | 1.62e-08 |
| $p$-value | 7.20e-05 | 3.97e-07 | 4.71e-01 | 2.29e-01 | 9.88e-01 | 1.00e+00 |
| CSA' | 1.79e-01 | 4.64e-02 | 1.60e-03 | 1.82e-03 | 9.17e-08 | 9.19e-12 |
| FDR CSA' | 3.55e-03 | 1.49e-01 | 8.91e-01 | 8.57e-01 | 9.96e-01 | 9.96e-01 |
| CSA$_\text{r}$ | 3.97e-01 | 5.07e-01 | 7.21e-02 | 1.20e-01 | 1.44e-03 | 1.27e-05 |
| FDR CSA$_\text{r}$ | 5.31e-03 | 2.06e-03 | 3.67e-01 | 1.69e-01 | 9.94e-01 | 9.96e-01 |
| $\chi^2_{\text{CSP}}$ | 8.69e+01 | 4.22e+01 | 1.00e+02 | 4.56e-01 | 5.63e+00 | 2.21e+01 |
| $p$-value | 1.17e-20* | 8.38e-11* | 1.23e-23* | 5.00e-01 | 1.76e-02 | 2.59e-06 |
| CSP | 3.44e+00 | 2.85e+00 | 3.90e+00 | 1.82e+00 | 3.91e+00 | 3.91e+00 |
| FDR CSP | 7.59e-03 | 1.58e-02 | 7.05e-03 | 3.93e-02 | 7.04e-03 | 7.04e-03 |

Table S2 Comparison of the $\chi^2$-test results and the FDRs corresponding to the values of CSA', CSA$_r$ and CSP for the six time samples taken under an symmetric MA for model B (see Fig. S9). The parameters values of the model are $c_{H_1} = c_{P_1} = c_{H_2} = c_{P_2} = 0$, $\beta = 0.00005$, $s = 0.6$, $\omega_1 = \omega_2 = 0.9$, $S_{1,\text{init}} = S_{2,\text{init}} = 41500$, $I_{11} = I_{12} = I_{21} = I_{22} = 4150$, $\delta_t = 0.001$, $b = 1$, $\gamma = 0.9$, and the sample sizes are $n_T = 200$ and $n_{\text{Inf}} = 100$. Asterisks (*) indicate significant p-values after Bonferroni correction for $10^7$ comparisons.

| | t=50 | t=55 | t=60 | t=250 | t=255 | t=260 |
|---|---|---|---|---|---|---|
| $\chi^2_{\text{CSA}}$ | 6.69e+01 | 6.69e+01 | 6.69e+01 | 6.69e+01 | 6.69e+01 | 6.69e+01 |
| $p$-value | 2.80e-16* | 2.80e-16* | 2.80e-16* | 2.80e-16* | 2.80e-16* | 2.80e-16* |
| CSA' | 8.18e-01 | 8.18e-01 | 8.18e-01 | 8.18e-01 | 8.18e-01 | 8.18e-01 |
| FDR CSA' | 0.00e+00 | 0.00e+00 | 0.00e+00 | 0.00e+00 | 0.00e+00 | 0.00e+00 |
| CSA$_\text{r}$ | 8.18e-01 | 8.18e-01 | 8.18e-01 | 8.18e-01 | 8.18e-01 | 8.18e-01 |
| FDR CSA$_\text{r}$ | 3.83e-04 | 3.83e-04 | 3.83e-04 | 3.83e-04 | 3.83e-04 | 3.83e-04 |
| $\chi^2_{\text{CSP}}$ | 4.18e-13 | 6.40e-15 | 2.93e-13 | 2.02e-14 | 1.19e-13 | 4.76e-13 |
| $p$-value | 1.00e+00 | 1.00e+00 | 1.00e+00 | 1.00e+00 | 1.00e+00 | 1.00e+00 |
| CSP | 6.47e-10 | 8.23e-12 | 6.04e-10 | 5.13e-10 | 2.93e-10 | 3.06e-10 |
| FDR CSP | 8.04e-01 | 8.04e-01 | 8.04e-01 | 8.04e-01 | 8.04e-01 | 8.04e-01 |

Table S3 Comparison of the $\chi^2$-test results and the FDRs corresponding to the values of CSA', CSA$_r$ and CSP for the six time samples taken under an GFG-interaction for model B (see Fig. S10). The parameters values of the model are $c_{H_1} = c_{P_1} = 0$, $c_{H_2} = c_{P_2} = 0.05$, $\beta = 0.00005$, $s = 0.6$,$\omega = 0.9$, $S_{1,\text{init}} = S_{2,\text{init}} = 41500$, $I_{11} = I_{12} = I_{21} = I_{22} = 4150$, $\delta_t = 0.001$, $b = 1$, $\gamma = 0.9$, and the sample sizes are $n_T = 200$ and $n_{\text{Inf}} = 100$.

| | t=50 | t=55 | t=60 | t=250 | t=255 | t=260 |
|---|---|---|---|---|---|---|
| $\chi^2_{\text{CSA}}$ | 8.12e-04 | 5.88e-04 | 4.50e-04 | 1.34e-04 | 1.37e-04 | 1.42e-04 |
| $p$-value | 9.77e-01 | 9.81e-01 | 9.83e-01 | 9.91e-01 | 9.91e-01 | 9.91e-01 |
| CSA' | 3.31e-05 | 2.44e-05 | 1.90e-05 | 4.77e-06 | 4.67e-06 | 4.56e-06 |
| FDR CSA' | 9.96e-01 | 9.96e-01 | 9.96e-01 | 9.96e-01 | 9.96e-01 | 9.96e-01 |
| CSA$_r$ | 2.85e-03 | 2.43e-03 | 2.12e-03 | 1.16e-03 | 1.17e-03 | 1.19e-03 |
| FDR CSA$_r$ | 9.90e-01 | 9.92e-01 | 9.93e-01 | 9.95e-01 | 9.95e-01 | 9.95e-01 |
| $\chi^2_{\text{CSP}}$ | 7.32e-03 | 8.26e-03 | 4.65e-03 | 5.99e-06 | 8.29e-06 | 1.11e-05 |
| $p$-value | 9.32e-01 | 9.28e-01 | 9.46e-01 | 9.98e-01 | 9.98e-01 | 9.97e-01 |
| CSP | 1.45e-02 | 1.62e-02 | 9.67e-03 | 3.35e-02 | 4.21e-02 | 5.16e-02 |
| FDR CSP | 8.04e-01 | 8.04e-01 | 8.04e-01 | 8.04e-01 | 7.94e-01 | 7.80e-01 |
